# Coalescent inference using serially sampled, high-throughput sequencing data from intra-host HIV infection

**DOI:** 10.1101/020552

**Authors:** Kevin Dialdestoro, Jonas Andreas Sibbesen, Lasse Maretty, Jayna Raghwani, Astrid Gall, Paul Kellam, Oliver G. Pybus, Jotun Hein, Paul A. Jenkins

## Abstract

Human immunodeficiency virus (HIV) is a rapidly evolving pathogen that causes chronic infections, so genetic diversity within a single infection can be very high. High-throughput “deep” sequencing can now measure this diversity in unprecedented detail, particularly since it can be performed at different timepoints during an infection, and this offers a potentially powerful way to infer the evolutionary dynamics of the intra-host viral population. However, population genomic inference from HIV sequence data is challenging because of high rates of mutation and recombination, rapid demographic changes, and ongoing selective pressures. In this paper we develop a new method for inference using HIV deep sequencing data using an approach based on importance sampling of ancestral recombination graphs under a multi-locus coalescent model. The approach further extends recent progress in the approximation of so-called *conditional sampling distributions*, a quantity of key interest when approximating coalescent likelihoods. The chief novelties of our method are that it is able to infer rates of recombination and mutation, as well as the effective population size, while handling sampling over different timepoints and missing data without extra computational difficulty. We apply our method to a dataset of HIV-1, in which several hundred sequences were obtained from an infected individual at seven timepoints over two years. We find mutation rate and effective population size estimates to be comparable to those produced by the software BEAST. Additionally, our method is able to produce local recombination rate estimates. The software underlying our method, Coalescenator, is freely available.

## Introduction

Human immunodeficiency virus (HIV) is a rapidly evolving pathogen that causes a chronic infection for an individual’s lifetime. As a consequence, the genetic diversity within a single infection can be very high. Important clinical variables, such as the rate of progression to AIDS and the set point viral load, are related to the diversity and evolution of the within-patient viral population (Shankarappa et al., 1999; Ross and Rodrigo, 2002; Williamson, 2003; Edwards et al., 2006; Lemey et al., 2007; Pybus and Rambaut, 2009), and so genetic data from these populations are of medical relevance in addition to providing insight into molecular evolutionary processes. However, population genomic inference from HIV sequence data can be challenging as result of high rates of mutation and recombination within a small RNA genome of approximately 10 kb. Furthermore, natural selection is expected to play an important role in shaping within-host HIV genetic diversity (Rouzine and Coffin, 1999; Neher and Leitner, 2010; Batorsky et al., 2011; Pennings et al., 2014). For example, phylogenies constructed from serially-sampled intrahost population sequences are typically characterized by a ladder-like topology (Shankarappa et al., 1999), indicating a rapid and continual turnover of genetic lineages as result of continual host immune selection. Although the relationship between census viral population size (i.e. viral loads) over the course of infection and virus kinetics is well understood for HIV (Nowak and May, 2000), relatively little is known about the effective population size during a single infection.

High coverage “deep” sequencing is now being used to address these problems and to investigate the genomic diversity of HIV, with several data sets already under study (Henn et al., 2012; Poon et al., 2012; Gall et al., 2013). For the study of within-host evolution of HIV patients, deep sequencing serves as a potentially powerful way to infer the evolutionary and ecological dynamics of the viral population at an unprecedented detail, particularly since sequencing can be performed at different timepoints during infection (Drummond et al., 2003). This is especially important for studying fast-evolving RNA viruses, where the substitution rates and effective population size may change through time. By utilizing the temporal structure of the sampled sequences, statistical power can greatly improve the precision of population demographic and evolutionary estimates (Pybus et al., 2000; Drummond et al., 2003). However, from the perspective of population genomic inference, deep sequencing data is unusual: high-throughput sequencing generates large numbers of short sequence reads, with each read representing 400–700 nt of a viral genome. Typically no virus particle is sampled twice, so that only a small fraction of the viral genome is sequenced for any given virion in the population. Genealogies can be estimated, for example, for each ∼500 nt sub-genomic partition, but each genealogy represents a different random sample of individuals from the same population. This presents peculiar challenges in genomic inference.

To infer parameters of an underlying evolutionary model from deep sequencing data, such as effective population size, mutation rate, and recombination rate, new theoretical and statistical approaches are therefore needed. In this article we work within a coalescent framework, and in particular its extensions to allow for serially-sampled, or *heterochronous*, sequences (Rodrigo and Felsenstein, 1999). (This is in contrast to the usual situation of *isochronous* sampling at a single fixed time.) The coalescent is a powerful and flexible framework for modelling the genealogy of a large, panmictic population, with many further extensions that incorporate changing population size, recombination, and recurrent mutations (see Hein et al., 2005, for a textbook introduction). It is a crucial component in the inference of the evolutionary dynamics of fast-evolving RNA viruses, which can be combined with epidemiological data in an approach known as *phylodynamics* (Grenfell et al., 2004). A potentially powerful method of inference under complicated coalescent evolutionary models is to proceed by computationally-intensive Monte Carlo simulation. The quantity of interest, such as the likelihood for the data, is estimated by averaging over a large number of the possible unobserved gene genealogies that could have given rise to the observed sequences. Genealogies of high posterior probability can be targeted by Markov Chain Monte Carlo (MCMC) or importance sampling (IS). There is now a sizeable literature on these and related approaches, focusing on various complications of the basic coalescent depending on the species under study. For example, there are methods to account for recombination [MCMC: Kuhner et al. (2000), Wang and Rannala (2008), Rasmussen et al. (2014); IS: Griffiths and Marjoram (1996), Fearnhead and Donnelly (2001), McVean et al. (2002), Griffiths et al. (2008), Jenkins and Griffiths (2011)], changing population size [MCMC: Beaumont (1999), Drummond et al. (2002, 2005), Wilson et al. (2003), Minin et al. (2008); IS: Griffiths and Tavar´e (1994a), Beaumont (2003), Leblois et al. (2014)], and heterochronous sequence data [MCMC: Drummond et al. (2002, 2005), Minin et al. (2008); IS: Beaumont (2003), Anderson (2005), Fearnhead (2008)].

However, for accurate inference using deep sequencing data from within-host virus populations we need to account for several of these complications simultaneously, and no existing methods are available for this task. In particular, current inference methods in this context have tended to omit the process of recombination (Pybus and Rambaut, 2009), limiting our understanding of the extent of the association between recombination and viral adaptation. However, the effective recombination rate is of an order of magnitude comparable to the mutation rate [for example, Shriner et al. (2004) found the recombination rate to be 5.5-fold greater than the mutation rate in HIV]; furthermore, recombination may play an important role in the evolution of drug resistance (Kellam and Larder, 1995; Neher and Leitner, 2010). There is also growing evidence that recombination rates are not constant along the genome (Fan et al., 2007; Archer et al., 2008). Thus, our goal in this paper is to develop a coalescent method of inference that can handle *all* of the following:

(i) Recombination,
(ii) High mutation rates,
(iii) Heterochronous sequences,
(iv) Missing data,
(v) Changing effective population size.

Here we take a novel importance sampling approach to tackle this problem. It is based on recent developments in the efficient computation of *conditional sampling distributions* [CSDs; Paul and Song (2010); Paul et al. (2011); see also Sheehan et al. (2013)]: the probability distribution of an additionally sampled haplotype, conditioned on the sampled haplotypes we have seen already. These are crucial in the design of efficient IS algorithms: Stephens and Donnelly (2000) noted an equivalence between designing an IS proposal distribution and approximating the (unknown) CSD. This work was later formalized by De Iorio and Griffiths (2004a), which subsequently allowed Griffiths et al. (2008) and Paul and Song (2010) to derive an accurate approximate CSD in the presence of crossover recombination. Finally, work by Paul et al. (2011) resulted in an efficient approximation of the CSD for multiple loci via a hidden Markov model (HMM).

Our work is most closely related to that of Griffiths et al. (2008), whose focus was a model able to account for recombination, but which was not designed with heterochronous deep sequencing data in mind, and thus exhibited only properties (i) and (ii) above. Extending this method to include (iii, iv, v) raises numerous methodological challenges that we address in further detail below. Briefly, we replace the CSD of Griffiths et al. (2008) with that of the sequentially Markov model of Paul et al. (2011). The latter is more efficient to compute and easily extended to blocks of completely linked sites. To allow for samples taken at different times we introduce an explicit time variable, which in turn determines when samples are inserted into the reconstruction of a genealogy backwards in time. Further, and in contrast to the imputation approaches of Fearnhead and Donnelly (2001) and Griffiths et al. (2008), our model allows for haplotypes to be only *partially* specified, assigning allelic states only at a subset of loci where possible. This dramatically reduces the state space of the model, and it also provides a convenient way of handling missing data.

Despite all of these contributions, application to large high-throughput datasets remains challenging, particularly for full-likelihood methods. In order to retain tractability, recent research has turned to approximations of the model, or of the likelihood, or both. The approach we take here is a principled method to find an accurate *full*-likelihood solution first by restricting attention to a *two-locus model.* Our two-locus model is then extended to multi-locus data by analysing selected pairs of loci separately and then aggregating these pairwise inferences, either by taking the median of the inferred estimates or by combining their likelihood surfaces via a pseudolikelihood [in particular, via a pairwise composite likelihood (McVean et al., 2002; Larribe and Fearnhead, 2011)]. As we describe below, we performed a simulation study in order to to demonstrate that our method can recover model parameters accurately.

The inferential process involves simulating ancestral trees backwards in time, and, in agreement with Griffiths and Tavar´e (1994b), Nielsen (1997), and Jasra et al. (2011), as we get closer to the MRCA, coalescence times increase greatly. This adds undesirably high variance to the inference, and extensive CPU time. To circumvent this, we further employ the *Time Machine* strategy developed by Jasra et al. (2011): stopping the simulation before the MRCA is reached whilst controlling for the bias; acquiring sizable computational saving and reduced variance.

To investigate the performance of our method on empirical data, we analysed HIV-1 RNA samples that had been extracted and sequenced at seven timepoints over a period of two years, from a patient enrolled in the control arm of the Short Pulse Antiretroviral Therapy at Seroconversion (SPARTAC) study. This patient received no anti-viral drugs. Nine regions from the whole HIV-1 genome alignments of these heterochronous data were then analysed using our model, and for comparison by the Bayesian MCMC coalescence approach implemented in BEAST (Drummond and Rambaut, 2007; Drummond et al., 2012). Both analyses provide strong agreement in the mutation rate, effective population size, and time to the most common recent ancestor. However, in addition, our approach also estimates recombination rates.

A C++ implementation of the algorithm is available from https://github.com/OSSCB2013 under the name `Coalescenator.` The program can process serially sampled, deep sequencing data from viral populations, to infer mutation rates, recombination rates, the effective population size, and times to recent ancestors.

## Materials and Methods

Model and notation. In this section we describe our notation and genealogical model. We begin by formulating a two-locus model; that is, given a pair of loci, call them A and B, recombination may occur at any position along the sequence separating them. We regard a locus as a fixed stretch of nucleotides, which may contain several polymorphic sites. Recombination within a locus is ignored.

*Demographic model.* Suppose we have data *D* = (*D*_-*K*_, *D*_-*K*+1_,…, *D*_0_) collected at time-points *t*_-*K*_ > *t*_-*K*+1_ > … > *t*_0_ = 0, with *t*_0_ being the most recent collection time and *t*_-1_, *t*_-2_,… extending into the past. We assume the effective population size (*N_e_*) to be constant between collection times, but it may change at these times. This piecewise-constant model is similar to that of Pybus et al. (2000). Except through their effects on *N_e_*, we otherwise ignore the effects of natural selection and population substructure.

To reduce the parameter space, viral load estimates at the sampling times can be used as a guide for constraining the relative magnitudes of the effective population sizes. In the simplest scenario (e.g. when viral load estimates are unavailable, unreliable, or we decide not to use them), a single, constant effective population size can be fitted, and in this paper we focus on the inference of a single effective population size parameter. In the results section, we investigate the robustness of our inference to this assumption.

*Mutation model.* Let *l_A_* and *l_B_* denote the lengths (in nt) of locus *A* and *B* respectively, and set *l* = *l_A_* + *l_B_* to be their total length. We assume that all nucleotides conform to a diallelic model, with alleles labelled arbitrarily as {0,1}. Each nucleotide mutates independently at the same rate and with symmetric transitions such that the mutation transition matrix for each nucleotide is 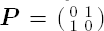. (In other words, *P_ij_* denotes the probability that, if the *i*th allele undergoes a mutation event, it changes to the *j*th allele.) Let 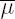 be the mutation rate per generation time per site, which we assume to be constant across sites, and let 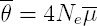 be the population-scaled mutation rate per generation per site when the effective population size is *N_e_.* Then 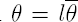 is the population-scaled mutation rate per generation time across both loci, with individual locus mutation rates given by 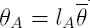 and 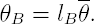. The overall mutation transition matrix for locus *ℓ* is

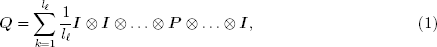
where ***I*** is the identity matrix and for each kth summand, ***P*** appears in the *k*th position in the direct product (Griffiths and Tavaré, 1994b). In other words, when a mutation occurs at locus *ℓ*, the resulting haplotype is chosen uniformly from amongst those differing from the parental haplotype at precisely one of the *l_ℓ_* sites. Because of the possibility of a high rate of mutation, this model allows for a site to have undergone more than one mutation event in its genealogical history. In practice we simulate mutation events only at polymorphic sites.

*Recombination model.* Suppose locus *A* and *B* are separated by a region of length *d* nucleotides, and assume a uniform recombination rate across this region. Let 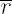 be the recombination rate per generation time per site, which we assume to be constant across sites. Therefore, if 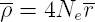 denotes the population-scaled recombination rate per generation time per site, then 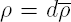 is the population-scaled recombination rate per generation time between locus *A* and *B.* We assume only crossover-type recombination, which implies an approximately linear relationship between short physical distance and recombination rate.

*Sample notation.* We sample *n* haplotypes at a given collection time; by *haplotype* we mean a sequence of nucleotides which we will later subdivide into a sequence of contiguous blocks of loci. Due to missing data, we observe *a* haplotypes specified only at locus *A, b* haplotypes specified only at locus *B*, and *c* haplotypes specified at both loci, so that *a* + *b* + *c* = *n.* Reads only partially overlapping with a locus are treated as missing at this locus. A haplotype (*i, j*) ∈ {0, 1}*^l_A_^* × {0, 1}*^l_B_^* is seen with multiplicity *c_ij_*, so that Σ_*i,j*_ *c_ij_* = *c.* Haplotypes with data missing at locus *A* or *B* are respectively denoted (*i*, *) and (*, *j*), with respective multiplicities *a_i_* and *b_j_*, and Γ_*i*_ *a_i_* = *a*, Γ_*j*_ *b_j_* = *b*. Let *c* = (*c_ij_*), ***a*** = (*a_i_*), and ***b*** = (*b_j_*); the complete dataset is thus written compactly as **n** = (***a, b, c***). As we reconstruct a genealogy for the sample backwards in time, recombination events create lineages each ancestral only at one of the two loci. The notation we have introduced to allow for missing data also allows us to leave unspecified the allelic types of these nonancestral recombinant loci: backwards in time, a recombination event replaces a type (*i,j*) with (*i*, *) and (*, *j*)

**Importance sampling.** Parameter estimation in population genetic models requires computation of the likelihood of the observed data *D* as a function of the model parameters Φ. We start by describing a method for estimating the likelihood *L*(Φ) for heterochronous data at two loci, with 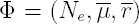. Our method also provides a weighted approximation to the posterior distribution of genealogical histories given this data, so it is straightforward to address questions of ancestral inference, such as the time to the most recent common ancestor (TMRCA) of the data. Note that under the standard (isochronous) coalescent model, it is not possible to identify mutation and recombination rates separately, as only their respective products with the effective population size can be identified. Serial sampling from a rapidly evolving population (relative to the spacing between samples) allows for the separate estimation of these three parameters, because a separate estimate of *N_e_* is then available. (To understand this, observe that with heterochronous data the times between sample collections appear in the likelihood expression, *L*(Φ). The timescale of the coalescent is governed by *N_e_*, which therefore now appears in *L*(Φ)—and so can be estimated—separately from its appearance inside the composite parameters 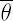 and 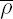. Further details are given by Rodrigo and Felsenstein, 1999, p259-262).

As it is not possible to write down *L*(Φ) = P(*D*; Φ) analytically, a *latent* genealogy variable, *G*, is introduced. The likelihood is then calculated by marginalising over *G:*

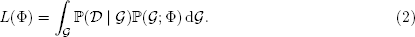

A naϊve Monte Carlo estimator for the integral in (2) is given by

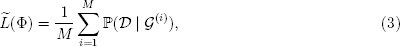

where *G*^(*i*)^ represent independent samples from P(*G*; Φ), the coalescent prior. This estimator has poor properties because P(*D* | *G*^(*i*)^ is 0 with high probability, resulting in 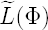 having very high variance (Stephens and Donnelly, 2000). Estimation of the integral (2) is therefore generally conducted using either MCMC or IS. The latter approach, which we follow here, was pioneered by Griffiths and Tavaré (1994b). An IS estimator is obtained by introducing an artificial *proposal distribution* ℚ and reweighting the samples; the estimator

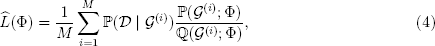

where *G*^(*i*)^ represent independent samples from ℚ(*G*; Φ), has lower variance than (3) provided ℚ(*G*; Φ) is chosen carefully. It can be shown that the *optimal* proposal distribution is the posterior P(*G* |*D*;Φ), in which case 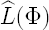 has variance 0 (Stephens and Donnelly, 2000). This distribution is unknown in general, but it can guide our intuition on how to choose ℚ(*G*; Φ).

Griffiths and Tavaré (1994b) noted that we can focus solely on genealogies compatible with the observed data by simulating them sequentially and by a reversal of time. To illustrate the idea we introduce some further notation similar to that of Stephens and Donnelly (2000):

First, regard the coalescent model as a Markov process forwards in time on unordered sets of haplotypes. The process starts at the most recent common ancestor (MRCA) of the sample, *H_-m_* ∈ {0, 1}^*l*^, and ends at an unordered set of *n* observed haplotypes, *H*_0_ ∈ ({0, 1}^*l*^)^*n*^, corresponding to the most recently observed data. The process visits a sequence of states *H* = (*H-_m_*, *H*_-*m*+1_,…, *H*_0_) as the genealogy is constructed forwards in time by mutations, coalescences, and recombinations; the known rates of these transitions are prescribed by the coalescent model. We refer to *H* as the *history* of the sample. The importance sampling estimator can then be decomposed as

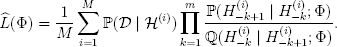

The numerators P(*H*_-*k*+1_ | *H*_-*k*_; Φ) are known from the coalescent model, and the denominators 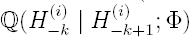 are specified by the proposal distribution. The key point is that this reformulation of the importance sampling procedure can be viewed as exploring the state space of latent histories *backwards* from the sample *H*_0_ to the MRCA *H_-m_.* One advantage of this is immediate: when the only data collection time is *t*_0_ = 0 then *D* = *H*_0_, and by choosing 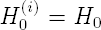 for each *i* = 1, …,*M* we can ensure that P(*D* | *H*^(*i*)^) = 1 for each *i*, so that no simulation is wasted. Moreover, the Markov specification of the proposal distribution helps to simplify our task of actually designing it.

A key observation of Stephens and Donnelly (2000) was that the posterior, conditional distribution of genealogies can be expressed in terms of the *conditional sampling distribution* (CSD), which is defined as the probability π[(*i*, *j*) | n; Φ] that a haplotype sampled from the population is of type (*i,j*) ∈ {0, 1}^*l*^ given that we have already sampled the haplotypes with configuration n. Although this distribution is as intractable as P(*G* | *D*;Φ), it provides a sensible starting point in the design of an efficient proposal distribution, since we have reduced the problem to one of finding a good approximation of π[(*i, j*) | n; Φ]. We emphasize that even if we use a proposal distribution based on an *approximation* to π[(*i, j*) | n; Φ] reweighting in (4) still provides us with an unbiased and consistent estimator.

**New proposal distribution.** Our aim then is to design a sequential proposal distribution ℚ(*H*_-*k*_ | *H*_-*k*+1_,; Φ), defining a Markov chain backwards in time which approximates P(*H* | *D*;Φ). Following the discussion of the previous section, to achieve this we use two steps: first, express the true backwards transition rates P(*H*_-*k*_ | *H*_-(*k*-1);_ Φ) in terms of the CSDs π[· | ·; Φ]; second, substitute the CSD for a well motivated approximation 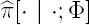. We address each of these steps in turn.

*Backwards transition rates.* For now, fix *N_e_* and assume a single sample collection time *t*_0_ = 0. Griffiths et al. (2008) obtained the backwards transition rates for a two-locus, finite-alleles model with isochronous samples taken from a population at equilibrium. Their formulation is efficient because it corresponds to ancestral recombination graphs (ARGs) in which only lineages carrying genetic material ancestral to the sample are simulated; entirely nonancestral lineages are integrated out. This dramatically reduces the state space and improves efficiency. However, the simulation continues to assign allelic types to the remaining nonancestral *loci.* For example, if a lineage in an ARG is ancestral to a member of the sample at locus *A* but not at locus *B*, then it is still necessary to assign an allelic status at locus *B.* In our application these loci represent tens or hundreds of nucleotides, and assigning a haplotypic status to such loci throughout the simulation is cumbersome. We therefore marginalize over these nonancestral loci too. The resulting backwards transition rates are given in Table A1, and the entries in this table are derived in Appendix A.

Thus far we have ignored changes in population size and heterochronous sampling. To account for these factors we introduce an explicit time variable *T* = (*T*_-*m*_, *T*_-*m* + 1_,…, *T*_0_ = 0), which contains the times between events in the history *H* (we now also include the collection of additional samples at times *t*_-1_, *t*_-2_, … as valid events). To calculate the likelihood, we now need to marginalise over both histories *H* and inter-event times *T:*

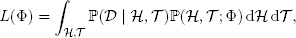
with IS estimator

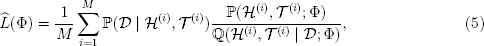
where the parameters of interest are 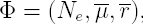, and (*H*^(*i*)^, *T*^(*i*)^) are independent samples from ℚ(*H*, *T* | *D*; Φ).

Our sequential proposal ℚ(*H*_-*k*-1_,*T*_-*k*-1_ | *H*_-*k*_,*T*_-*k*_;Φ) is described below. Suppose our genealogical reconstruction has proceeded backward a time 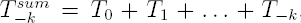, past the sample collection time *t*_-*j*_. We first sample an event time backwards according to an exponential distribution with rate depending on the current sample size (*a,b,c*) and the current effective population size *N*_-*j*_. In apprehension of the appearance of new samples at given times in the past, we measure time in chronological units rather than generation units (Rodrigo and Felsenstein, 1999). Then the waiting time *T* (in days) for the next event, is exponentially distributed with density

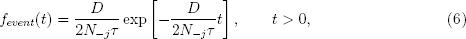
where *D* = *n*(*n* - 1) + (*a* + *c*) *θ_A_* + (*b* + *c*) *θ_B_* + *ρc* is (twice) the total prior rate of events and *τ* is the generation time in days. (The parameters *θ_A_*, *θ_B_*, and *ρ* are redefined each time the effective population size changes; e.g. 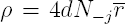) We sample a time *T* from (6): if the corresponding event time is more recent than the next (pastward) sample collection time i.e. 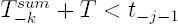, then we accept this time and define our proposal by

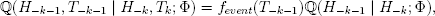
with ℚ(*H*_-*k*-1_ | *H*_-*k*_; Φ) defined as in Table A1—only we replace each instance of π[· | ·; Φ] with an approximation 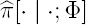 developed below. Otherwise, with probability

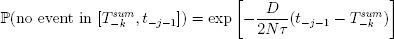
the next event is the insertion of additional samples at the next collection time, and we set the next waiting time to be 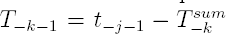. At collection time *t*_-*j*-1_, we insert the samples *D*_-*j*-1_ collected at time *t*_-*j*-1_ to our current configuration *H*_-*k*._ That is, if *D*_-*j*-1_ = *m* and *H_-k_* = n then we set *H*_-*k*-1_ = *m* + n. We then update the effective population size parameter to N_-j_-1, and continue this iterative procedure until we have both passed the oldest collection time and reached a MRCA for each locus. The overall procedure is illustrated in Figure 1.

**Figure 1:**
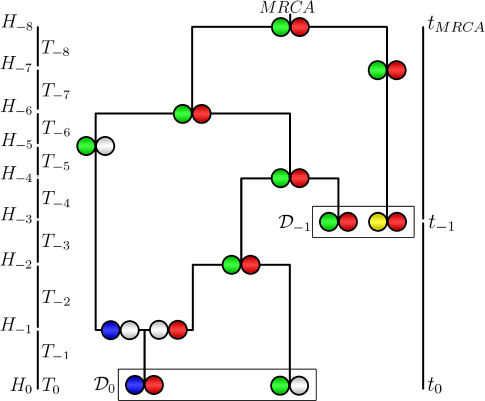
Illustration of our two-locus recombination model: a sampled history *H* and inter-event times *T.* The two loci of each haplotype are each represented by a circle. White circles represent an unspecified locus and colored circles indicate the allelic type at that locus. For example *H_0_* consists of types (blue, red) and (green, *). There are two sampling times and the collected samples are represented by the leaves of the tree (marked by rectangles). Time is measured in chronological units and run backwards from the most recent collection time, *t*_0_ = 0, to the most recent common ancestor *t_MRCA_.* Ancestral lineages are represented by black lines. At a coalescence event, two lineages are joined together; the model allows coalescence between fully-specified haplotypes (*H*_-4_), between a fully-specified and partially-specified haplotype (*H*_-6_), and between two partially-specified haplotypes (*H*_-2_). At a recombination event, two lineages are created and their haplotypes are partially specified: one of the two loci becomes non-ancestral and its allele type is left unspecified (*H*_-1_). At the next collection time *t*_-1_, a new sample *D*_-1_ is added to the existing lineages *H*_-2_: *H*_-3_ = *H*_-2_ ∪ *D*_-1_; and the effective population size is allowed to change.

*Conditional sampling distribution.* Our proposal distribution is obtained by substituting an approximate CSD 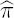 for π in Table A1. We use the the HMM approach devised by Paul et al. (2011), yielding an algorithm linear in both the number of loci and the number of haplotypes (i.e. linear in sequence length and sample size, respectively). It is a relatively accurate approximation of the true CSD and practical to compute. Briefly, the new haplotype (*i,j*) is sampled given an existing configuration n by assuming that the genealogy for the latter is a simple, improper “trunk” ancestry: one extending infinitely far back into the past with no mutations, recombinations, or coalescences (see Figure 2 and Paul and Song, 2010). The alleles of the new haplotype are determined by allowing its ancestral lineage to undergo mutation and recombination, and to coalesce into this trunk ancestry at appropriate rates. This framework can be cast as an HMM across loci, whose emissions are the observed alleles at the newly sampled haplotype, and whose hidden state is the pair *s_ℓ_* = (*τ_ℓ_,i*), where *τ_ℓ_* is the absorption time of this locus into the trunk, *i* ∈ {0,1}^*l_ℓ_*^ is the type of the lineage into which absorption occurs (Figure 2), and *l_ℓ_* is the length of this locus (in nt).

Although this framework is obviously a strong simplification of the true underlying genealogical process, the resulting CSD 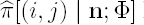 has many sensible properties (Paul et al., 2011). However, this approach does not lend itself immediately to handling unspecified alleles. We therefore make several further modifications to the CSD of Paul et al. (2011) in order to construct a practicable IS algorithm, as we now describe.

*Emission probabilities.* For our model, where all nucleotides are diallelic and mutate independently, we can simplify the emission probabilities used in the HMM calculation— increasing the computational efficiency greatly. Given a hidden state *s_ℓ_* = (*τ_ℓ_, i*) of the Markov chain at locus *ℓ*, the emission probability mass function for observed state *j* ∈ {0, 1}^*l_ℓ_*^ is given by:

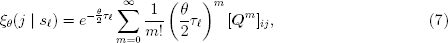
where ***Q*** is the mutation transition matrix (1). Because sites within a locus mutate independently in our model, we can reformulate this as

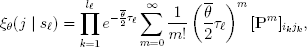
where we recall that 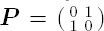. We can simplify this even further:

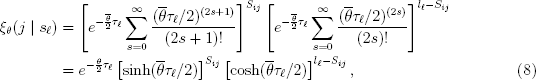
where *S_ij_* denotes the number of nucleotide differences between *i* and *j.* Thus, we have successfully eliminated the infinite sum in the emission distribution (7). Because the hidden state *τ_ℓ_* is continuous, we follow Paul et al. (2011) and employ Gaussian quadrature to construct a discrete HMM. We chose Laguerre-Gauss quadrature (Abramowitz and Stegun, 1972, Table 25.9) with 16 points. Applying the forward algorithm to this HMM yields the required CSD. The expression for the emission distribution in this discretized model can also be reduced to a closed-form formula (Appendix B).

*Emission probability when the absorbing hidden state is unspecified.* The method of Paul et al. (2011) does not deal with the case where the locus of interest at the absorbing hidden state is unspecified, i.e. calculating emissions of the form *ξ_θ_* (*j* | (*τ_ℓ_*, ∗)). In a two-locus model this occurs, for example, when the absorbing state is the haplotype (*i*, ∗) for some *i* and we are interested in the emission distribution at locus B. In this case, we *condition* on choosing an absorbing haplotype with the allele at this locus specified (Figure 3(B)). When there are no haplotypes in the trunk ancestry for which the locus of interest has a specified allele, we sample the allele from the stationary distribution of ***Q*** [eq. (1)]; that is, we pick uniformly from all possible 2^*l_ℓ_*^ haplotypes at this locus:

**Figure 2.**
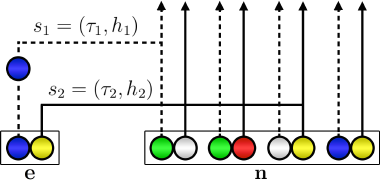
Illustration of the sequential interpretation for a realization of 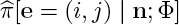 for two loci. The dotted and full lines, respectively represent the marginal genealogies (*S*_1_, *S*_2_) at locus *A* and *B.* The hidden state at locus *A* is *s*_1_ = (*τ*_1_, *h*_1_). Haplotype *h*_1_ would carry a green allele at its first locus, but a mutation results in the observed blue allele. The hidden state at locus *B* is *s*_2_ = (*τ*_2_, *h*_2_). *h*_2_ carries a yellow allele at its second locus, and no mutation occurs on the marginal genealogy at this locus. If there is no recombination, *s*_2_ = *s*_1_, but here a recombination occurs before *τ*_2_ and the absorption time for the second locus is *τ*_2_ ≠ *τ*_1_. As in Figure 1, white circles represent loci with unspecified alleles.

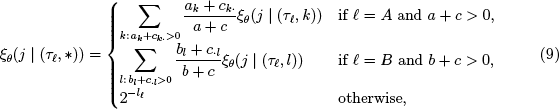
where the trunk ancestry has configuration **n** = (**a, b, c**) and we have used the notation *c_k_.* = Σ_*l*_ *l_kl_*, *c._l_* = Σ_*k*_ *c_kl_*.

*Emission probability for partially observed haplotypes.* If the observed haplotype has an unspecified locus (i.e. we seek to calculate 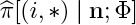 or 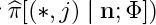, then we transform this to a one-locus problem with a degenerate HMM summing over hidden states at a single locus (Figure 3(**A**)). If the absorbing hidden state is unspecified (Figure 3(**B**)) then we choose uniformly from the informative trunk lineages as described above.

*CSD for recombination event.* To model recombination, we require π[{(*i*, ∗), (∗, *j*)} | **n**; Φ], the CSD for two partially observed haplotypes (see Table A1). This quantity satisfies the following decompositions and symmetries:

**Figure 3:**
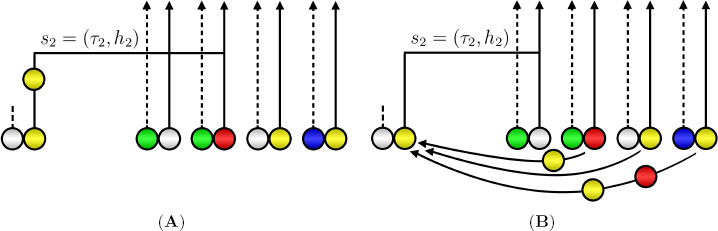
Sampling 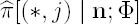, with the observed allele *j* represented by a yellow circle. (**A**): Absorption at the second lineage of the trunk ancestry for which the second locus is specified (red allele). Mutation event is still allowed in this one-locus model, as illustrated here by a mutation from a red to yellow allele. (**B**): Absorption at the first lineage of the trunk ancestry for which the second lineage is unspecified. In such cases we choose uniformly from the other informative lineages as the absorbing state.

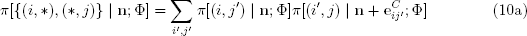

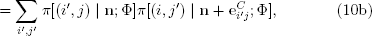

and

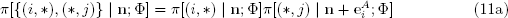

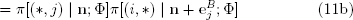

To estimate 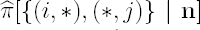, Griffiths et al. (2008) use relationship (10) by substituting approximate CSDs 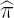 for each fully-observed haplotype. In addition, they noted that 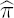 may not satisfy symmetries (10a,b)—and so they take the average of the two expressions. However, this strategy in averaging over all the alleles at the unspecified loci is very computationally intensive. Here, we instead use relationship (11), and substitute the approximate CSDs 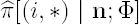 and 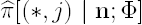 from the previous section (*Emission probability for partially observed haplotypes*). We still take the average of the two expressions (11a,b) to account for asymmetries.

*Forward Transitions.* To summarize, solving the forward equation associated with the HMM described above allows us to compute an approximate CSD, 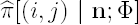, for each (*i, j*) ∈ {0,1}^*l*^ as well as for the special cases (*i*, ∗) and (∗, *j*). Plugging this into Table A1 defines an IS proposal distribution for sampling genealogical histories associated with the two loci. Equation (5) then provides an unbiased estimator of the likelihood. We have not yet specified how to compute the numerator in (5); to do this we also need the prior probabilities P(*H, T*; Φ). These can be computed easily and sequentially using the rightmost column of Table A1 (with minor modifications to account for the simulation of *T*). Table A1 covers events involving coalescence, mutation, and recombination. In addition, we also need to calculate the probability of subsampling the observed haplotypes *D_-j_* from among the lineages in the reconstructed genealogy, for each collection time *t_-j_.* Thus, there is a contribution to the numerator of (5) given by the hypergeometric distribution (Beaumont, 2003): if *H_-k_* = **n**∗ is the configuration at the collection time (including the additional samples) and *H*_-*k*_ + 1 = **n** is the configuration just before (more recently than) the addition of these samples, then

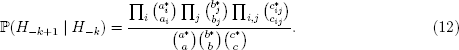

After decomposing P(*H, T*; Φ) in this manner it remains to compute the final term P(*H*_-*m*_) for the probability of the type of the MRCA. Under our uniform mutation model, the stationary distribution of the mutation process and thus the type of the MRCA is uniformly distributed such that *P*(*H_-m_*) = 2^-*l*^. Finally, the probability of the data given a history, *P*(*D* | *H,T*), is simple an indicator for whether the configurations at the leaves of the simulated history coincide with the observed data. By construction this is identically equal to 1.

**Further algorithmic improvements.** In this section, we present two modifications to our model for reducing the computational overhead, as well as two strategies for extending the two-locus model to process multi-locus data.

*Proposal distribution.* The work of Stephens and Donnelly (2000), De Iorio and Griffiths (2004a), Griffiths et al. (2008), Paul and Song (2010), and Paul et al. (2011) suggests that our approach to IS will have attractive statistical efficiency. However, generating proposals according to Table A1 requires the evaluation of all possible events for all haplotypes in the current configuration *H_-k_*, which may be too costly for complex samples. Instead we therefore make a simple modification, similar to the one suggested by Fearnhead and Donnelly (2001), which is a further approximation to the proposal suggested by Table A1: Consider the waiting time to the next event as the minimum of independent competing exponential times for the events involving each of the (*i,j*),(*i*,∗), and (∗,*j*) haplotypes. Now sample a haplotype to be involved in the next event according to the *prior* probability of its involvement. In this calculation we exclude the possibility of a coalescence between dissimilar haplotypes, resulting in the probabilities given in Table A2. Next, given the chosen haplotype we choose the event it is involved in with probabilities proportional to the relevant rows in Table A1; now only those rows need to be computed.

*Time machine.* As reported by Griffiths and Tavaré (1994b), Nielsen (1997), and Jasra et al. (2011), as we simulate the tree backwards and closer to the MRCA, the simulation times increase greatly. This long simulation run results in undesirably high variance of the likelihood estimate, and extensive CPU time—a particular drawback of an IS approach. To circumvent this, Jasra et al. (2011) considered stopping the simulation before the MRCA is reached, in an approach termed the *Time Machine.* The bias from this action is then characterized. First, it depends on the underlying mixing of the evolutionary process: the closer to the root node of the tree, the process is able to ‘forget’ its initial condition. Secondly, it depends on the specific distribution of the process at the stopping (exit) time. Using the IS algorithm of Stephens and Donnelly (2000), they investigated the bias-variance effect of stopping simulations at the first time back that the number of lineages had decreased to 1%, 2%, 5%, 10%, 25%, and 50% of the original sample size *n.* They concluded that stopping at 5% strikes the right balance between computational efficiency and numerical accuracy.

We conservatively adopt their approach: stopping at *T*_*MRCA*-5_, the first time the backwards simulation reaches five ancestors. At this exit time, the alleles of the remaining lineages are drawn from a uniform distribution. Indeed, we experience sizable computational saving and reduced variance, without noticeable effect on the quality of the estimates.

*Estimation of local and global parameters.* The method described above estimates the likelihood *L*(Φ) for heterochronous deep sequencing data at a pair of loci. Recall that each locus is a stretch of nucleotides within which recombination is *ignored*, but between which recombination is *explicitly modelled.* By searching for Φ that maximizes *L*(Φ), we obtain the *pairwise maximum likelihood estimate (pairwise MLE)* for the mean mutation rate 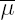 and effective population size across the pair, and the mean recombination rate 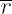 between the pair.

This two-locus IS algorithm can be used to analyse multi-locus data. Given a region of length *κ* (typically ∼ 400-500 nt), we partition it into loci of length *δ* (typically ∼ 50 nt), defining a collection of *ω* = *κ*/*δ* loci. We run the IS algorithm on pairs of non-adjacent loci {(*i,j*): |*j* - *i*| > 1; *i,j* ∈ {1, 2,…, *ω*}}. Adjacent pairs are excluded because recombination between loci separated by a single nucleotide is expected to be negligible. The resulting pairwise MLEs are then indicative of how the local population parameters are distributed within the region.

In practice, the size for the partitioned loci, *δ*, would be chosen based on the read length distribution of the data, so that a large proportion of the sample at any particular locus will be fully specified and thus can be included in our analysis. (Recall that if a sequence contains even a few missing nucleotides at a given locus the rest of the information at this locus is discarded and we consider it as a block of missing data.) A balance should be struck: smaller locus length allows inference of finer recombination and mutation rate variation across the region, at the expense of higher computational complexity.

It is also of interest to have a single, global estimate for the population parameters that is representative of the whole region. An ideal situation is a short region (typically ∼ 500 nt) where the evolutionary behavior is relatively homogeneous. In such cases, one can take the *median* or *mean* of pairwise MLEs as a simple global estimator; we found the median to be more robust. An alternative, more sophisticated global estimator is via a *pairwise composite likelihood* (reviewed in Larribe and Fearnhead, 2011), which we include for comparison with the median of pairwise MLEs. In this strategy, the two-locus IS algorithm is run on each pair *i,j* ∈ {1, 2,…, *ω*} of loci such that 1 < |*i* — *j*| ≤ Δ, for some threshold distance Δ. The global parameters are inferred by maximizing pairwise composite likelihood:

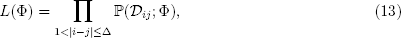
where *D_ij_* denotes the marginal data restricted to the pair of loci *i* and *j.* Equation (13) is similar to the composite likelihood of McVean et al. (2002), using pairs of multi-nucleotide loci rather than pairs of single-nucleotide polymorphisms (SNPs), as suggested by Jenkins and Griffiths (2011).

**Figure 4:**
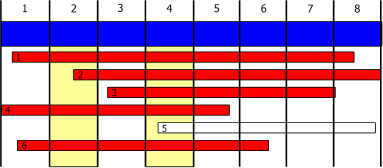
Estimation of the local and global parameter estimates using the two-locus IS algorithm. A region of interest, typically ∼ 4–500 nt, is shown as a blue segment. This region is partitioned into smaller loci, e.g. 50 nt. Sequence reads are shown as thinner horizontal bars. For non-adjacent pairs of loci, the two-locus engine computes pairwise MLEs as local estimates of the population parameters. Here, a pairwise comparison between locus 2 and 4 is illustrated (yellow shading). Reads that fully cover at least one of the two loci are highlighted in red and are used for the inference: three complete haplotypes (reads 1, 4, and 6) and two partial haplotypes (reads 2 and 3). These pairwise inferences can be combined to reach the global parameter for the whole region. Two approaches are described in the text: by taking the median of the pairwise MLEs or via a pairwise composite likelihood.

Introducing a threshold Δ is attractive for both statistical and computational reasons (Larribe and Fearnhead, 2011), and there are several additional reasons why Δ ought to be small in our application. First, the short-read nature of the data means that if |Δ*i* — *j*| > Δ, then many samples will be missing at one or both of the two loci. Second, since our loci represent blocks of nucleotides rather than single SNPs, parameter variation along the sequence implies we should concentrate on proximate pairs of loci. Third, our modelling of complete blocks of nucleotides instead of isolated SNPs prohibits the pre-computation of an exhaustive list of pairwise likelihoods, as is possible in McVean et al. (2004) for example. For these reasons we focus on the composite likelihood (13) in which Δ = 2.

These procedures are illustrated in Figure 4.

**Simulated data.** Simulated heterochronous datasets were generated using the software package **NetRecodon** (Arenas and Posada, 2010), using a Jukes-Cantor substitution model (Jukes and Cantor, 1969) and with each nucleotide frequency value equal to 0.25. Population parameters were chosen to be comprable to those previously described for HIV: 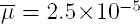 per site per generation, 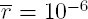 per site per generation, *N_e_* = 10^3^, *τ* = 1.8 days (e.g. Shriner et al., 2004; Batorsky et al., 2011; Neher and Leitner, 2010). Each simulation was set to produce *n_c_* sequences of length *κ* = 500 nt, at four collection times separated by 252 days, totaling *n* = 4*n_c_* sequences.

Under the above conditions, four datasets were generated under slightly different scenarios. The first pair of datasets, called *Const-120* and *Const-600*, were generated under a constant population size. *Const-120* had *n_c_* = 30 reads sampled at each collection time, totaling *n* = 120 reads. *Const-600* had a 5-fold increased sample size to *n_c_* = 120 reads at each collection time, totaling *n* = 600 reads. The second pair of datasets, *Dynamic-120*, and *Dynamic-600*, had the same population parameters and samples sizes as *Const-120* and *600-const* respectively. However, they were generated under a fluctuating viral population such that *N_e_* corresponded to 1,000, 2,000, 4,000, and 500 at the four collection times (from the most recent to the earliest). This models a population undergoing both an expansion and a bottleneck through time.

**Real data.** The subject studied here was enrolled in the control (no therapy) arm of the Short Pulse Antiretroviral Therapy at Seroconversion (SPARTAC) study. EDTA plasma samples were obtained at the start of the study (at an estimated 12 days from HIV-1 seroconversion) and at 28, 120, 176, 373, 429 and 695 days after the start of the study. Viral RNA was extracted, whole HIV-1 genomes were sequenced in a pool of 21 samples on 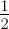 PicoTiterPlate using a Genome Sequencer FLX Titanium XL instrument (Roche/454 Life Sciences), and a consensus sequence for each time point was derived as previously described (Gall et al., 2012). The seven consensus sequences were aligned using MAFFT v6.857 (Katoh et al., 2002), and a consensus sequence of this alignment was used as a reference sequence for mapping of all reads, with suffixes representing the time points, using Burrows-Wheeler Aligner v0.5.9 (Li and Durbin, 2010). The resulting SAM file was converted into a FASTA file using a custom Java script. The distribution of read lengths and positions across the genome are given in Table S_1_ and Table S_2_ respectively.

*Accession numbers for sequencing data.* The Roche/454 Life Sciences sequencing data obtained in this study is available from the EMBL/NCBI/DDBJ Sequence Read Archive sample accession numbers ERS661087-ERS661093.

*HIV genome analysis.* Nine regions were selected from the whole HIV genome alignment of mapped reads, each region approximately 600 nt in length and comprising samples from all time points. Furthermore, we ensured that these regions were in-frame and non-overlapping, spanning the four main open-reading frames (i.e. gag, pol, env, and nef). Reads covering at least 95% of a region were retained in our analysis. The average number of sequences per time point was 34 (range 20-65). Further details about the sequence dataset for each region can be found in Table S3.

*BEAST analysis.* To evaluate the performance of Coalescenator in estimating population parameters of interest, primarily the effective population size, *N_e_*, and mutation rate, 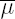, we re-analysed the sequence alignments for the nine regions described above using the Bayesian MCMC coalescent approach implemented in BEAST (v1.8.0 Drummond and Rambaut, 2007; Drummond et al., 2012). Specifically, a codon-structured nucleotide substitution model (Shapiro et al., 2006), a constant size population demographic model, and a strict molecular clock prior were employed. Two independent MCMC chains of 50 million steps were run to assess convergence and adequate mixing.

## Results

**Simulated data.** To evaluate the validity of our model implementation, Coalescenator, we simulated heterochronous datasets under known population parameters. Employing this set, we assessed Coalescenator’s performance with respect to the number of Monte Carlo iterations, the sample sizes of the data, and the data’s underlying population size dynamics. For each study, we searched through a three-dimensional parameter space of 11 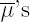, 11 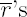, and 6 N_e_’s that best explain the observed diallelic sites in the data (see Table S4 for the summary of the sites classification and the numbers of sites used). We assessed whether parameter combinations with the highest likelihoods corresponded to the true parameters that generated the data. 11×11×6 = 726 parameter combinations of the following were used:

- 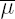 1 × 10^-7^, 1 × 10^-6^, (1,2.5,5,7.5) × 10^-5^, (1,2.5,5,7.5) × 10^-4^, 1 × 10^-3^.
- 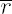1 × 10^-8^, (1,2.5,5,7.5) × 10^-7^, (1,2.5,5,7.5) × 10^-6^, 1 × 10^-5^, 1 × 10^-4^.
- *N_e_*: 250, 750, 1000, 1250, 1750, 3000.

*Performance for data from constant population size.* The behavior of Coalescenator when analysing data generated under constant population size was assessed using datasets *Const-120* and *Const-600* (see Materials and Methods). Coalescenator was run with a constant population size model.

First, the performance of local inference using pairwise MLEs was assessed by analysing a single pair of loci of lengths *l_A_* = *l_B_* = 50 nt at positions (1, 50) and (101, 150). Figure 5 shows the likelihood surfaces for *Const-120*, with 100 and 500 Monte Carlo iterations respectively; a further increase to 1000 and 5000 iterations is shown in Figure S_1_. Here, across various choices of *N_e_*, there is a clear clustering of higher likelihoods near the true parameter values. With only 100 iterations, the pairwise MLE recovers the correct mutation rate (among the 11 parameter values considered), and estimates the recombination rate at 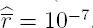. The MLE for the effective population size is further off-target, at 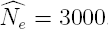. By increasing the number of iterations to 500 [Figure 5(B)], there was a marked improvement in the likelihood surface; it is now much more peaked, and the recombination rate estimate improves to within a factor of two of its true value (although for this experiment the effective population size is still overestimated). Overall, parameter estimates are accurate for 500 iterations, and certainly to within an order of magnitude. For this dataset, we also investigated a more ambitious increase to 1000 and 5000 Monte Carlo iterations (Figure s_1_); the recombination rate estimate sees modest further improvement, but at a cost of infeasible computing requirements should we hope to scale this to real datasets. Indeed, it is worth emphasising that these experiments use relatively few Monte Carlo iterations; typical IS approaches to coalescent inference might use several orders of magnitude more. Our aim here is to stretch our algorithm when only a few iterations are available, so that it can feasibly be scaled up to complex multi-locus datasets. The results presented here suggest that the sophistication of our algorithm adequately compensates for the dearth of CPU time, and comparisons with longer runs show that even as few as 100 iterations give decent estimates. The nature of the data, with its high mutation rates and heterochronicity, may also assist in this exploration of tree space.

Figure 6 shows the corresponding likelihood surfaces for *Const-600.* The likelihood hypersurface concentrates in a similar manner with respect to the true parameter values, but this time there is a shift towards the correct effective population size. This suggests that Coalescenator performs favorably with increasing sample size.

It is also worth commenting on the general shape of the likelihood hypersurfaces in Figures 5 and 6. One might expect to see a ridge corresponding to a negative correlation in the estimation of mutation and recombination rates, as alternative ways of resolving any incompatibilities in the data. In fact the main ridge, evident from several panels in the figures, is parallel to the recombination axis. This reflects the simple fact that recombination is inherently more difficult than mutation to estimate to a given accuracy. A second, weaker ridge is evident in the contours along lines of constant *N_e_µ*, as might be expected. That no other ‘ridges’ seem to be discernible from these likelihood hypersurfaces is encouraging, as it is consistent with the fact that the data contains sufficient information for mutation and recombination parameters to be estimated separately (and that our method can reconstruct the likelihood surface to sufficient accuracy).

The above analysis shows that by using the information from a single pair of loci of 50 nt each, Coalescenator is able to arrive at a sensible indication of the population parameters for the whole 500 nt region. This raises two questions:

1. For effective inference, how far apart should the two loci be?
2. A pairwise MLE provides only partial information since it utilises a subset of the whole region. How then, should one come up with the global estimate for the whole region?

To answer the first question, inference quality was assessed with respect to the separation *d* of a pair of loci. The 500 nt sequence region was divided into 10 neighboring loci of 50 nt, and for each of the 36 pairs of non-adjacent loci, pairwise MLEs for the population parameters were calculated. Figure *S*_2_ shows how the MLEs vary with *d.* We observed that the mutation rate estimate is stable across separation distance. However, this is not the case with the recombination estimates. Specifically, (i) Variance in MLEs for fixed *d* is greater than for mutation; and (ii) With increasing *d*, recombination rate estimates (per site) seem to be lower.

These two observations could have been anticipated to some extent; the signal for recombination in genetic data is generally weaker than that of mutation. Furthermore, with increasing separation between two loci the signal for recombination can become ’saturated’, so that recombination events are undetected and recombination rates are underestimated. More precisely, the curvature of the likelihood curve for *ρ* is flatter when *ρ* is greater (Chan et al., 2012). Ultimately, this provides the basis in deciding the appropriate constraint on locus pair separation for reliable inference. This is about 50-100 nt, and hereonwards we proceeded by computing pairwise MLEs for pairs of loci separated by 50 nt.

To address the second challenge of obtaining global parameter estimates, we assessed both of the two approaches discussed in the section *Estimation of local and global parameters.* The first approach is to take the median of the pairwise MLEs. Figure 7 shows pairwise MLEs and the corresponding median across the datasets. We see that the median can effectively capture the true estimates, albeit with a slight overestimation of *N_e_.* There is also greater variance in estimates for the recombination rates, which in turn reflect the confidence we should have in these estimates.

The second approach is via a pairwise composite likelihood (13), in which likelihood surfaces for pairs of loci separated by 50 nt are multiplicatively combined. Figure S3 shows the composite likelihood surfaces for dataset *Const-120.* The MLEs using 100 and 500 Monte Carlo iterations are both of high accuracy, and differ only slightly in the recombination rate estimates; this further illustrates the fast convergence of Coalescenator. With 500 iterations, the true mutation and recombination rates are captured, although the effective population size is off-target at 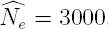. Figure S4 shows the result for *Const-600*, where the larger sample size also provides accurate estimates of the effective population size. For example, with 500 iterations the joint MLE is 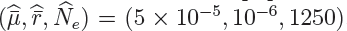, only two cells away from the true values (on this grid). These estimates are also included in Figure 7 for direct visual comparison with the first approach.

**Figure 5:**
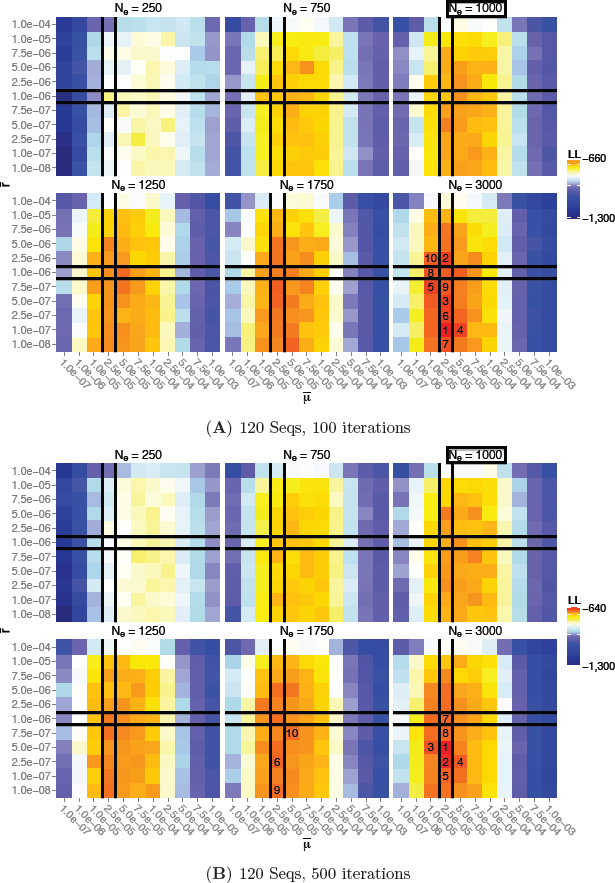
Likelihood surfaces on a pair of loci at positions (1,50), (101,150), for dataset *Const-120*, using 100 and 500 Monte Carlo iterations. Cells correspond to the searched parameters, colored by log-likelihoods, with the top 10 estimates numbered. The true mutation, recombination, and population parameters, are: 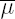 = 2.5 x 10^-5^, 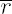 = 10^-6^, *N_e_* = 10^3^.

**Figure 6:**
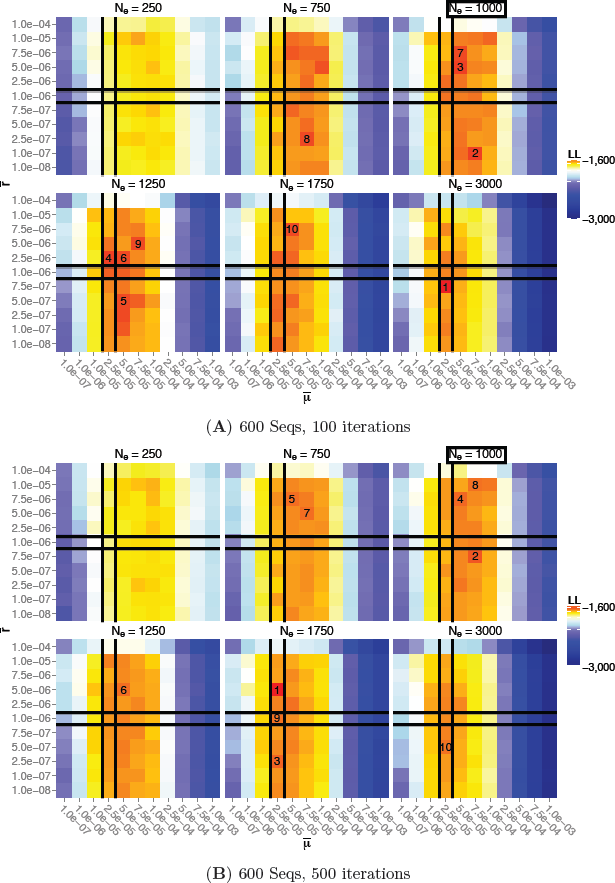
As in Figure 5 but for dataset Const-600.

**Figure 7:**
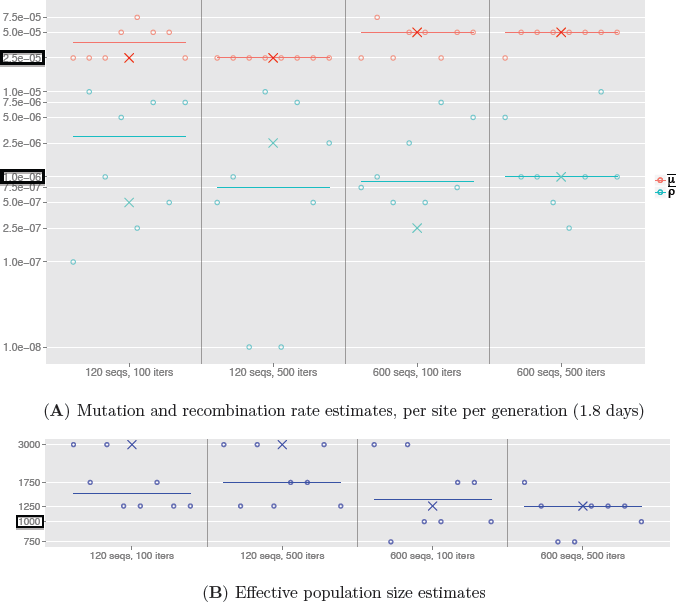
Population parameter estimates for simulated datasets generated under a constant population size model. The effect of using different combinations of sample size and number of Monte Carlo iterations is compared. Circles correspond to the pairwise MLEs between neighbouring pairs of loci (those separated by 50 nt). The horizontal lines correspond to the median of the pairwise MLEs. Crosses indicate the pairwise composite likelihood estimates.

*Performance for data from dynamic population sizes.* The behavior of Coalescenator for analysing data generated under dynamic population sizes was assessed using datasets *Dynamic-120* and *Dynamic-600.* Figure 8 shows the pairwise MLEs and the global estimates based on the median and pairwise composite likelihoods, for both datasets. Each estimate was based on 100 Monte Carlo iterations.

With the incorrect assumption of a constant population size, `Coalescenator` slightly overestimates the mutation rates. The pairwise MLEs for recombination rates increase in accuracy for the larger sample size, but there is greater variability compared to the analysis in the previous section on *Const-120* and *Const-600.* Effective population size estimates are still of very reasonable accuracy and stability. This suggests that `Coalescenator` is reasonably robust to unmodelled changes in population size, and can be reliably used in this way. Of course, it is straightforward to extend our algorithm to impose population size changes between sample collection times, perhaps guided by viral load information, and even to *infer* changes in effective population size. However, in the latter case we found that our chosen ranges of sample size and Monte Carlo iterations were insufficient to compensate for the significant increase in model complexity.

As all the cases here and the previous section illustrate (Figures 7 and 8), the two methods for providing global parameter estimates—taking the median of pairwise MLEs and constructing a composite likelihood estimate—have generally shown good agreement, which corroborate each other’s accuracy. The former seems to give similar or better global measures than the latter, which seems slightly more susceptible to outlier estimates, so we focus on the median of pairwise MLEs to provide our global summaries hereonwards. Clearly, strategies more sophisticated than a simple median might be able to aggregate pairwise estimates more efficiently, and we leave this for future work.

*Variance of the importance Weights.* For each of the four simulated datasets, we assessed the stability of Coalescenator’s likelihood estimates by computing a coefficient of variation 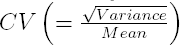 of the importance weights [i.e. the summands in (5)]. We used 100 Monte Carlo iterations for each particular parameter combination, analysing each pair of loci separated by 50 nt. There are 11x11x6 = 726 parameter combinations and 8 pairs of loci, resulting in 5, 808 *CVs* being computed for each dataset.

Table 1 summarises the distribution of the coefficient of variation for the four simulated datasets. The variance appears to be well constrained, with the majority of *CVs* under 0.001. The larger *CVs* originate from parameter combinations that deviate furthest from the ground truth parameters. While a small *CV* (equivalently, a large effective sample size) is not a guarantee of good performance, it is consistent with the idea that the proposal distribution is not too skewed in its exploration of the space of ARGs.

*Running Time.* `Coalescenator` scales approximately linearly with respect to the sample size and the number of Monte Carlo iterations. For dataset *Const-120*, a likelihood calculation for a single parameter combination using 100 iterations takes on average 10 seconds when run in parallel on a 16-core computer with a 2.0 GHz processor. For dataset *Const-600*, the corresponding calculation increases 5-fold to about 50 seconds.

**Real data analysis.** `Coalescenator` was run on nine HIV regions from HIV genome alignments collected at seven time points from an HIV infected patient over a period of two years (see MATERIALS and METHODS). The sequence data is summarised in Table S3.

Each gene region is 600 nt long, but some reads had missing data, chiefly up to the first 30 nt and the last 30 nt of the region. Although **Coalescenator** can handle missing data, for the purpose of this analysis we concentrate on the central positions (51, 550) of each gene region. Sites containing ambiguous nucleotides, indels, and non-diallelic sites were filtered out. Table S5 summarises the sites classification, as well as the total and breakdown of the number of sites used for each gene region.

**Figure 8.**
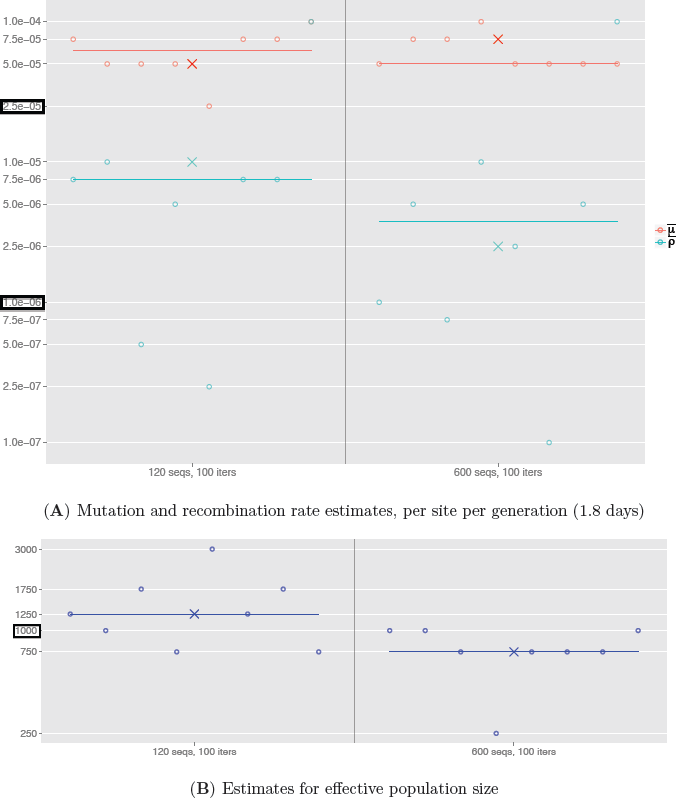
Population parameter estimates for simulated datasets generated under a dynamic population size model. Coalescenator was run under a constant population model, and this analysis shows its robustness to unmodelled changes in population size. Circles correspond to the pairwise MLEs between neighboring pairs of loci (those separated by 50 nt). The horizontal lines correspond to the median of the pairwise MLEs. Crosses indicate the pairwise composite likelihood estimates.

**TABLE 1:** Assessment of the variation of the importance weights, under different simulated datasets. For each dataset, a *CV* of the importance weights was calculated for each run of 100 Monte Carlo iterations on a particular parameter combination and a pair of loci separated by 50 nt. This results in 5, 808 *CV*s being computed in total for each dataset. Each cell in the table shows the proportion of these *CV*s that are below the indicated thresholds.

| *CV Threshold:* | < 0.15 | < 0.1 | < 0.05 | < 0.01 | < 0.001 |
| --- | --- | --- | --- | --- | --- |
| *Const-120* | 1 | 1 | 0.92 | 0.83 | 0.83 |
| *Const-600* | 1 | 0.86 | 0.86 | 0.86 | 0.86 |
| *Dynamic-120* | 1 | 1 | 0.99 | 0.95 | 0.95 |
| *Dynamic-600* | 1 | 0.91 | 0.9 | 0.88 | 0.88 |

The parameter combinations extended those used in the simulation study. Additional gridpoints (2.5,5,7.5) × 10^-5^ for the recombination rate were considered, to account for observation in a preliminary run, that several MLEs for recombination rates across the regions seem to be between 1 × 10^-5^ and 1 × 10^-4^. Following the simulation study, each gene region is partitioned into ten 50 nt loci, and likelihood surfaces were generated for pairs of loci separated by 50 nt nt using 100 iterations.

Figure 9 shows the evolutionary parameter estimates across each region. The median of the pairwise MLEs and the pairwise composite-likelihood estimates agree quite closely for the mutation and effective population size estimates, and all lie in the range [2.5 × 10^-5^,1 × 10^-4^] for mutation rate estimates in each region. There was greater variability across the genome for effective population size estimates: for example, all median-based estimates were in the range 875-1000 in regions encompassing env, while the median-based estimate for the region pol 2836-3436 had 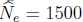. There was some disagreement between the two estimates of recombination rate. The pairwise composite likelihood estimates seem to be less stable; its estimate in the region env 6415-7015 is 2-10 times lower than any other region, and one tenth of the estimate obtained using the median of pairwise MLEs. (As discussed above, this behavior was also evident in the simulation study.) We prefer therefore to use the median of the pairwise MLEs as our regional summaries, with the uncertainty expressed by the variability in individual MLEs.

**Figure 9:**
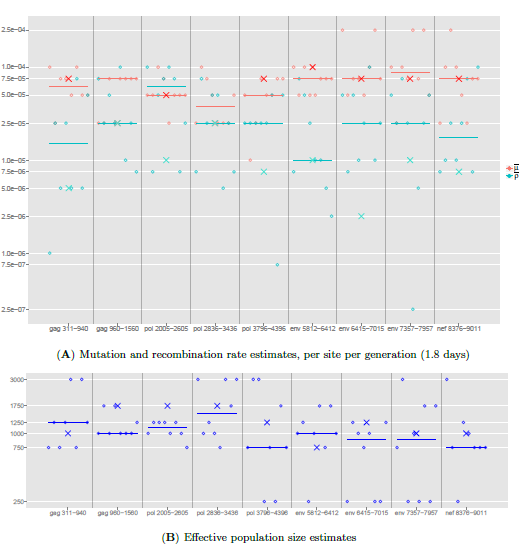
Population parameter estimates for nine HIV gene regions. Data is from HIV genome alignments collected at seven time points over the period of two years. Circles correspond to the pairwise MLEs between neighbouring pairs of loci separated by 50 nt. The horizontal lines correspond to the median of the pairwise MLEs. Crosses indicate the pairwise composite likelihood estimates.

It is worth noting that some pairwise MLEs are suggestive of a higher mutation rate in parts of the env region (as described previously by Alizon and Fraser, 2013). There is also some variability in the median-based recombination rate estimates across regions, but this is consistent with Monte Carlo sampling error. A single pairwise likelihood surface between loci (1, 50) and (101, 151) and a pairwise composite likelihood surface is shown in Figure 10 for the env 6415-7015 region. This serves as a sanity check that we indeed have convergence to a plausible likelihood surface, since each gridpoint is simulated independently.

**Figure 10:**
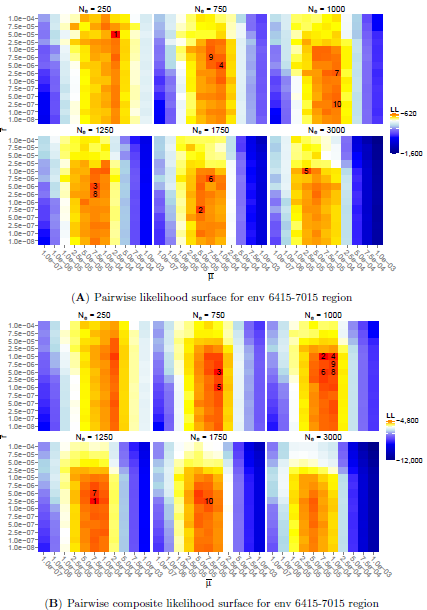
Likelihood surfaces for the env 6415–7015 region, using 100 Monte Carlo iterations. Cells correspond to the searched parameters, colored by log-likelihoods, with the top 10 estimates numbered. (**A**) Likelihood surface for a single pair of loci within the region. (**B**) Pairwise composite likelihood aggregating all valid pairs of loci within the region.

Finally, we compared our estimates against the output of the corresponding analysis using **BEAST** (Drummond and Rambaut, 2007; Drummond et al., 2012) (we do not compare recombination rates—recall that **BEAST** is unable to infer these. Our recombination rate estimates are summarised separately in Table 2.) The comparison is illustrated in Figure 11. The raw values for all the estimates are reported in detail in Table S6. The mutation 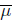 and recombination 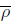 rates estimates from **Coalescenator** were converted into the number of substitutions and recombinations per site per year, respectively. *N_e_* denotes effective population size, and the time to the most recent common ancestor *T_M RCA_*, or the time until we reach 5 ancestors *T*_*M RCA*-5_, is given in years. Estimates of mutation rates and *T_M RCA_*s show very good agreement, though our method seems to be more cautious in estimating variation in *N_e_* across regions. Across the mutation rate estimates, the only notable differences are our higher estimates for the second gag region and the first pol region (each greater by a factor of ≈ 4). This discrepancy could reflect that **Coalescenator** can be prone to slight overestimation of mutation rates, as suggested by the simulation results on *Dynamic-120* and *Dynamic-600.*

**Figure 11:**
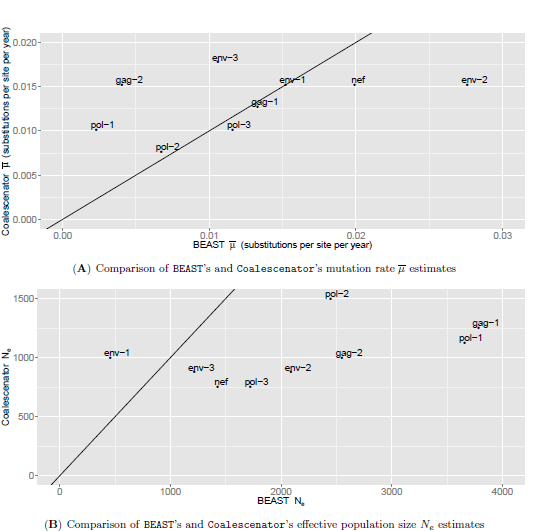
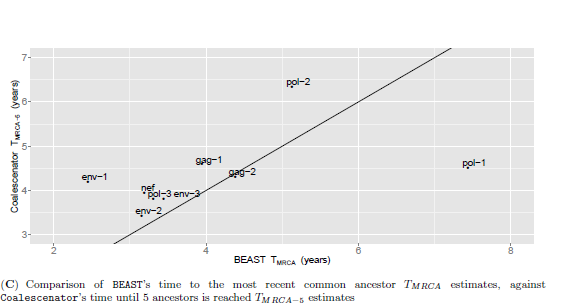
Comparison of **BEAST’s** parameter estimates against **Coalescenator’s**. An identity line *y* = *x* is plotted in each case, to visually assess the agreement of the estimates by the two programs. The following abbreviations are used for the nine HIV gene regions: gag-1 = gag 311-940, gag-2 = gag 960-1560, pol-1 = pol 2005-2605, pol-2 = pol 2836-3436, pol-3 = pol 3796-4396, env-1 = env 5812-6412, env-2 = env 6415-7015, env-3 = env 7357-7957, nef = nef 8376-9011. (**A**) Comparison of mutation rate 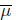 estimates, converted to the number of substitutions per site per year. (**B**) Comparison of effective population size *N_e_* estimates. (**C**) Comparison of **BEAST’s** time to the most recent common ancestor *T_M RCA_* estimates, against Coalescenator’s time until 5 ancestors is reached *T*_*M RCA*-5_ estimates, given in years.

**TABLE 2:** **Coalescenator’s** recombination rate 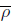 estimates, given in the number of recombinations per site per year.

| Region | $\bar{\rho}$ |
| --- | --- |
| gag 311–940 | $3.04 \times 10^{-3}$ |
| gag 960–1560 | $5.07 \times 10^{-3}$ |
| pol 2005–2605 | $1.27 \times 10^{-2}$ |
| pol 2836–3436 | $5.07 \times 10^{-3}$ |
| pol 3796–4396 | $5.07 \times 10^{-3}$ |
| env 5812–6412 | $2.03 \times 10^{-3}$ |
| env 6415–7015 | $5.07 \times 10^{-3}$ |
| env 7357–7957 | $5.07 \times 10^{-3}$ |
| nef 8376–9011 | $3.55 \times 10^{-3}$ |

## Discussion

We have presented a method capable of handling many of the challenges associated with modelling evolution of rapidly evolving viral populations measured using high-throughput sequencing. It is based on recent developments in the approximation of *conditional sampling distributions* (CSDs) (Stephens and Donnelly, 2000; De Iorio and Griffiths, 2004a; Griffiths et al., 2008; Paul and Song, 2010; Paul et al., 2011), and the adoption of this work into a practical IS algorithm should be a useful contribution in itself. The model we derive handles recombination by allowing for partially specified haplotypes and thus avoids the inflation of the model state-space caused by alternative imputation procedures. It also provides a natural way of dealing with missing data. One limitation is that data is lost when we simply consider an entire locus as unobserved if any nucleotides are missing; another possibility for future research would be to impute short stretches of missing nucleotides within a locus.

High-throughput sequence data from viral genomes presents many challenges. We have focused in particular on handling high mutation rates, recombination, heterochronous sampling, and missing data. Each of these presents its own hurdles; dealing with them simultaneously raises numerous further complications. In this paper we have therefore also explored several statistical and algorithmic techniques for reducing the large computational overhead. In particular, employing the *Time Machine* strategy developed by Jasra et al. (2011) proves to be effective in saving computational time, whilst controlling the bias and variance. Nevertheless, the overhead remains large, and it is likely that further implementation strategies will have to be explored in future as sample sizes continue to grow. For example, one advantage of an IS approach is that it is especially suited for parallisation using the rapidly growing GPU processing (Lee et al., 2010). Lee et al. focused on MCMC and IS as case studies, and concluded that significant performance boost can be achieved for IS using GPU, whereas **BEAST’s** MCMC is inherently less suitable for exploitation using GPU’s massive parallelisation.

In spite of these challenges, it is encouraging that our new method performs comparably to **BEAST** in the analysis of a deep sequencing dataset generated from a serially sampled acute HIV infection. Like previous studies based on coalescent inference, **BEAST** and **Coalescenator** also indicate that the effective population size is of the order ∼ 10^3^ (Brown, 1997). While there has been great interest in measuring the effective population size of within-host HIV infections, especially in regards to addressing whether evolution is largely stochastic or deterministic (Rouzine et al., 2014), it is important to note that estimates of effective population size from methods based on the coalescent framework do not typically consider natural selection or other factors that may lead to variation in offspring distribution. Consequently, in populations where variation in reproductive success is likely to be present, such as in within-host HIV infections, interpretation of the effective population size is confounded. Instead, a more accurate interpretation of effective population size is that it represents a measure of population turnover or relative genetic diversity. We further note that in this paper we have focused our analyses on inference of a single effective population size parameter, while a priority for future work will be also to make inference of temporal changes in this parameter computationally tractable.

The estimates of recombination rate from **Coalescenator** strongly agree with non-coalescent based estimates obtained by Neher and Leitner (2010) and Batorsky et al. (2011). This is encouraging as the sequence data analysed in this study has been sampled from an early phase of infection, in contrast to previous studies that utilised data from patients with a much longer follow-up (Neher and Leitner, 2010; Batorsky et al., 2011). Consequently, this demonstrates the applicability of our method even in spite of limited accumulation of diversity in the viral population.

Despite the complicated evolutionary model considered in this paper, we have omitted other mechanisms that are likely to be important to within-host HIV evolution. Our simplified model of population size changes provides a restrictive model for the phases of exponential expansion and contraction HIV and other viral populations are known to go through (Nowak and May, 2000). A natural solution to this problem would be to fit exponential functions between consecutive time-points. More importantly however, selective responses to the host immune system are known to be an important evolutionary factor (Edwards et al., 2006; Lemey et al., 2006), having an effect on intrahost genealogies that is not fully captured merely by adjusting the effective population size. Other commonly used methods such as **BEAST** (Drummond et al., 2002, 2005; Minin et al., 2008; Drummond and Rambaut, 2007; Drummond et al., 2012) also ignore the effects of selection on genealogies (as well as ignoring recombination); thus, developing methods that can robustly account for both selection and recombination (and indeed other factors) remains a challenging task. Another important consideration is population substructure: HIV is known to undergo *compartmentalization*, forming distinct subpopulations at different anatomical sites and in different cell types (Ewing et al., 2004). Incorporating substructure and migration into IS algorithms is in principle straightforward and has been achieved in other applications (Bahlo and Griffiths, 2000; De Iorio and Griffiths, 2004b; Griffiths et al., 2008), though it obviously introduces yet more computational burden.

Exploring more complicated mutation models would also be interesting, though the diallelic model we consider here allows for a simple analytical solution, which in turn allows for more efficient computation. However, it should be straightforward to allow for time-varying mutation rates as a proxy for the effects of selection. Inclusion of indels into the mutation model would also be desirable; inspection of the HIV data (cf. Table S5) shows that these are highly prevalent. The model also considers only cross-over recombination; however, McVean et al. (2002) suggested that a gene-conversion model may be appropriate for bacteria and viruses. Incorporating this type of recombination would also be a worthwhile extension.

Finally, we remark that in our analysis we tuned our algorithm to analyse data generated under the 454 platform. However, it should be straightforward to adapt our work for other platforms such as Illumina paired-end sequencing, in which a haplotype would now comprise a pair of reads. Since the number of reads generated by this method is typically several-fold larger, this raises further computational and statistical challenges which form the basis of ongoing work.

## Acknowledgements

This research was initiated during the 2013 Oxford Summer School in Computational Biology. We gratefully acknowledge Sarah Fidler, Steve Kaye, Jonathan Weber, Myra McClure, and the SPARTAC Trial participants and investigators. The work was supported by the Wellcome Trust and the NIHR Biomedical Research Centre funding scheme at Imperial College Healthcare NHS Trust and University College London Hospitals NHS Foundation Trust. We would also like to acknowledge funding from the Novo Nordisk Foundation (JAS, LM) and the EPSRC (PAJ, Grant EP/L018497/1). JR is supported by the Oxford Martin School. OGP received funding from the European Research Council under the European Union’s Seventh Framework Programme (FP7/2007-2013) / ERC grant agreement no. 614725-PATHPHYLODYN. We thank James Anderson, Farah Colchester, Pierre Haas, Thomas Leitner, István Miklós, and Yee Whye Teh for helpful comments on this work. The authors would like to acknowledge the use of the University of Oxford Advanced Research Computing (ARC) facility in carrying out this work (http://dx.doi.org/10.5281/zenodo.22558).

## Appendix

## A Backwards Transition Rates in the Arg

The distribution on the most recent event back in time in a two-locus ARG, *conditioned* on observing the sample configuration *H*-_*k*_ = **n**, is given in Table A1. This distribution is expressed in terms of the (unknown) CSD *π*[(*i,j*)| **n**; Φ] and transitions are marginalized over nonancestral genetic material; thus for example a recombination event changes a (*i,j*) haplotype to a (*i*, ∗) and a (∗, *j*), with ‘∗’ denoting unspecified alleles. [Such a marginaliza-tion approach was proposed by Jenkins and Griffiths (2011), but was based instead on an infinite sites model]. The corresponding forwards transition probabilities are also given in the rightmost column of Table A1. Here, e_*i*_ denotes a unit vector with *i*th entry 1 and the rest 0, with e_*ij*_ denoting a unit matrix whose only nonzero entry is a 1 at position (*i*,*j*). The notation 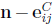 is shorthand for (***a, b,c*** - **e***_ij_*), and so on, and we write *c*_*i*·_ = Σ_*j*_ *c_ij_* and *c*_·*j*_ = Σ_*i*_ *c_ij_* for the marginal counts of fully specified haplotypes.

**TABLE A1:** Possible backwards transitions from state *H*_-*k*-1_ = **n** to state *H*_-*k*-1_ in a two-locus ARG in which loci are permitted to remain unspecified. Backward rates are expressed in terms of the CSD *π*[· | ·; Φ]. e_*i*_ denotes a unit vector with *i*th entry 1 and the rest 0, with e_*ij*_ denoting a unit matrix whose only nonzero entry is a 1 at position (*i*, *j*). The notation 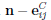 is shorthand for (***a, b, c*** − e_*ij*_), and so on, and we write *c_i_* = Σ_*j*_ *c_ij_* and *c*_·*j*_ = Σ_*i*_ *c_ij_* for the marginal counts of fully specified haplotypes.. The normalizing constant *D* denotes (twice) the total prior rate of events, *D* = *n*(*n* - 1) + (*a* + *c*)*θ_A_* + (*b* + *c*)*θ_B_* + *ρc*

| Event | $H_{-k-1}$ | $\mathbb{P}(H_{-k-1} \mid H_{-k})$ | $\mathbb{P}(H_{-k} \mid H_{-k-1})$ |
| --- | --- | --- | --- |
| Coalescence of two $(i, j)$s | $\mathbf{n} - \mathbf{e}_{ij}^C$ | $\frac{c_{ij}(c_{ij}-1)}{D} \cdot \frac{1}{\pi[(i, j) \mid \mathbf{n} - \mathbf{e}_{ij}^C; \Phi]}$ | $\frac{c(c_{ij}-1)}{D}$ |
| Coalescence of $(i, *)$ with $(i, *)$ or $(i, j)$ | $\mathbf{n} - \mathbf{e}_i^A$ | $\frac{a_i(a_i + 2c_i - 1)}{D} \cdot \frac{1}{\pi[(i, *) \mid \mathbf{n} - \mathbf{e}_i^A; \Phi]}$ | $\frac{a(a_i + 2c_i - 1)}{D}$ |
| Coalescence of $(*, j)$ with $(*, j)$ or $(i, j)$ | $\mathbf{n} - \mathbf{e}_j^B$ | $\frac{b_j(b_j + 2c_j - 1)}{D} \cdot \frac{1}{\pi[(*, j) \mid \mathbf{n} - \mathbf{e}_j^B; \Phi]}$ | $\frac{b(b_j + 2c_j - 1)}{D}$ |
| Coalescence of $(i, *)$ and $(*, j)$ | $\mathbf{n} - \mathbf{e}_i^A - \mathbf{e}_j^B + \mathbf{e}_{ij}^C$ | $\frac{2a_i b_j}{D} \cdot \frac{\pi[\{(i, *), (*, j)\} \mid \mathbf{n} - \mathbf{e}_i^A - \mathbf{e}_j^B; \Phi]}{\pi[(i, j) \mid \mathbf{n} - \mathbf{e}_i^A - \mathbf{e}_j^B; \Phi]}$ | $\frac{2ab(c_{ij} + 1)}{(c + 1)D}$ |
| Mutation of $(i, j)$ to $(k, j)$ | $\mathbf{n} - \mathbf{e}_{ij}^C + \mathbf{e}_{kj}^C$ | $\frac{\theta_A P_{ki}^A c_{ij}}{D} \cdot \frac{\pi[(k, j) \mid \mathbf{n} - \mathbf{e}_{ij}^C; \Phi]}{\pi[(i, j) \mid \mathbf{n} - \mathbf{e}_{ij}^C; \Phi]}$ | $\frac{\theta_A P_{ki}^A (c_{kj} + 1)}{D}$ |
| Mutation of $(i, j)$ to $(i, l)$ | $\mathbf{n} - \mathbf{e}_{ij}^C + \mathbf{e}_{il}^C$ | $\frac{\theta_B P_{lj}^B c_{ij}}{D} \cdot \frac{\pi[(i, l) \mid \mathbf{n} - \mathbf{e}_{ij}^C; \Phi]}{\pi[(i, j) \mid \mathbf{n} - \mathbf{e}_{ij}^C; \Phi]}$ | $\frac{\theta_B P_{lj}^B (c_{il} + 1)}{D}$ |
| Mutation of $(i, *)$ to $(k, *)$ | $\mathbf{n} - \mathbf{e}_i^A + \mathbf{e}_k^A$ | $\frac{\theta_A P_{ki}^A a_i}{D} \cdot \frac{\pi[(k, *) \mid \mathbf{n} - \mathbf{e}_i^A; \Phi]}{\pi[(i, *) \mid \mathbf{n} - \mathbf{e}_i^A; \Phi]}$ | $\frac{\theta_A P_{ki}^A (a_k + 1)}{D}$ |
| Mutation of $(*, j)$ to $(*, l)$ | $\mathbf{n} - \mathbf{e}_j^B + \mathbf{e}_l^B$ | $\frac{\theta_B P_{lj}^B b_j}{D} \cdot \frac{\pi[(*, l) \mid \mathbf{n} - \mathbf{e}_j^B; \Phi]}{\pi[(*, j) \mid \mathbf{n} - \mathbf{e}_j^B; \Phi]}$ | $\frac{\theta_B P_{lj}^B (b_l + 1)}{D}$ |
| Recombination of $(i, j)$ | $\mathbf{n} - \mathbf{e}_{ij}^C + \mathbf{e}_i^A + \mathbf{e}_j^B$ | $\frac{\rho c_{ij}}{D} \cdot \frac{\pi[\{(i, *), (*, j)\} \mid \mathbf{n} - \mathbf{e}_{ij}^C; \Phi]}{\pi[(i, j) \mid \mathbf{n} - \mathbf{e}_{ij}^C; \Phi]}$ | $\frac{\rho c(a_i + 1)(b_j + 1)}{(a + 1)(b + 1)D}$ |

To derive the entries in this table we use sampling exchangeability and Bayes’ theorem (see Stephens and Donnelly, 2000; De Iorio and Griffiths, 2004a). For example, to obtain the first row of Table A1, with 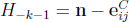, note that sampling exchangeability implies

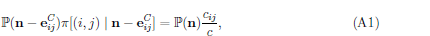

since both sides are equal to the probability that we sample the configuration **n** and then subsample a type (*i*, *j*); on the left by selecting the fully specified haplotype we sampled most recently, and on the right by subsampling from ***c***. [Our convention for ‘unordered’ is that P(**n**) = *σ*(***a, b, c***) · P(*A_n_*), where *σ*(***a, b, c***) = *a*!*b*!*c*!/(П*_i_a_i_*! П*_j_b_j_*! П*_i,j_ c_ij_*!) and *A_n_* denotes any *particular* ordering of the sample configuration **n**. In this convention, whether or not a haplotype is fully specified is not considered to be ‘random’ that is, the missingness or otherwise of either allele need not be assigned a probability. Instead, we can think of this as having *conditioned* on the missingness status of each locus.]

Now, by Bayes’ theorem,

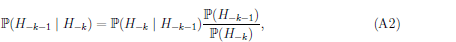

with

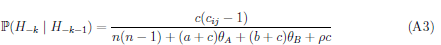
known from the coalescent prior. Probabilities such as (A3) can be obtained, for example, from the recursive system of equations studied by Ethier and Griffiths (1990) and Jenkins and Song (2009) (for example, multiply eq. (5) of Jenkins and Song (2009) by *σ*(***a,b,c***), rearrange to express the system in terms of P(**n**), and then read off the coefficients of the resulting system).

The entry for P(*H*_-*k*-1_ | H_-*k*_) in the first row of Table A1 follows by combining (A1), (A2), and (A3). The remaining entries follow similarly. At the bottom row, *π*[{(*i*, ∗), (∗, *j*)} | 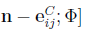 denotes

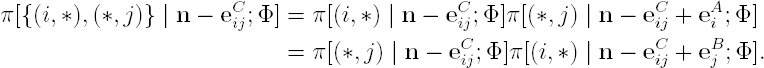

**TABLE A2:** Proposal probability of choosing a particular (*i*,*j*), (*i*, *), or (*, *j*), to be involved in the next event back in time, for each *i*,*j*. These probabilities are given up to a normalizing constant found by summing all these events over *i* and *j.*

| Haplotype | Probabilities up to a normalizing constant |
| --- | --- |
| $(i, j)$ | $c_{ij}(c_{ij} - 1 + \theta + \rho + a_i + b_j)$ |
| $(i, *)$ | $a_i(a_i - 1 + b + c_i + \theta_A)$ |
| $(*, j)$ | $b_j(b_j - 1 + a + c_j + \theta_B)$ |

## B Emission Probabilities for Gaussian Quadrature

After applying Gaussian quadrature it is necessary to compute discretized emission proba bilities of the form

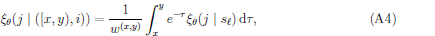

where 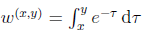 (Paul et al., 2011, eq. 17). In light of (8), we are also able to reduce this to a closed-form formula as follows. Substituting (8) into (A4), we find

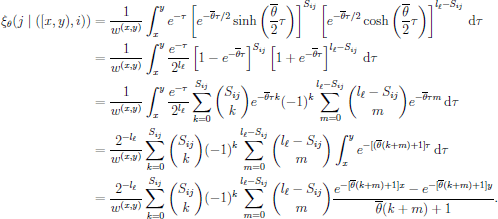

## References

Abramowitz, M. and Stegun, I., editors. Handbook of mathematical functions with formulas, graphs, and mathematical tables. Tenth printing, volume 55 of National Bureau of Standards Applied Mathematics Series. United States Department of Commerce, 1972.

Alizon, S. and Fraser, C. (2013). Within-host and between-host evolutionary rates across the HIV-1 genome. Retrovirology, 10 (1) 49.

Anderson, E. C. (2005). An efficient Monte Carlo method for estimating *N_e_* from temporally spaced samples using a coalescent-based likelihood. Genetics, 170, 955–967.

Archer, J., Pinney, J. W., Fan, J., Simon-Loriere, E., Arts, E. J., Negroni, M., and Robertson, D. L. (2008). Identifying the important HIV-1 recombination breakpoints. PLoS Computational Biology, 4, e10000178.

Arenas, M. and Posada, D. (2010). Coalescent simulation of intracodon recombination. Genetics, 184, 429–437.

Bahlo, M. and Griffiths, R. (2000). Inference from gene trees in a subdivided population. Theoretical Population Biology, 57, 79–95.

Batorsky, R., Kearney, M. F., Palmer, S. E., Maldarelli, F., Rouzine, I. M., and Coffin, J. M. (2011). Estimate of effective recombination rate and average selection coefficient for HIV in chronic infection. Proceedings of the National Academy of Sciences, 108, 5661–5666.

Beaumont, M. A. (1999). Detecting population expansion and decline using microsatellites. Genetics, 153, 2013–2029.

Beaumont, M. A. (2003). Estimation of population growth or decline in genetically monitored populations. Genetics, 164, 1139–1160.

Brown, A. J. L. (1997). Analysis of HIV-1 env gene sequences reveals evidence for a low effective number in the viral population. Proceedings of the National Academy of Sciences, 94, 1862–1865.

Chan, A. H., Jenkins, P. A., and Song, Y. S. (2012). Genome-wide fine-scale recombination rate variation in Drosophila melanogaster. PLoS Genetics, 8, e1003090.

De Iorio, M. and Griffiths, R. C. (2004a). Importance sampling on coalescent histories I. Advances in Applied Probability, 36, 417–433.

De Iorio, M. and Griffiths, R. C. (2004b). Importance sampling on coalescent histories II. Advances in Applied Probability, 36, 434–454.

Drummond, A. J. and Rambaut, A. (2007). BEAST: Bayesian evolutionary analysis by sampling trees. BMC Evolutionary Biology, 7, 214.

Drummond, A. J., Nicholls, G. K., Rodrigo, A. G., and Solomon, W. (2002). Estimating mutation parameters, population history and genealogy simultaneously from temporally spaced sequence data. Genetics, 161, 1307–1320.

Drummond, A. J., Pybus, O. G., Rambaut, A., Forsberg, R., and Rodrigo, A. G. (2003). Measurably evolving populations. Trends in Ecology & Evolution, 18, 481–488.

Drummond, A. J., Rambaut, A., Shapiro, B., and Pybus, O. G. (2005). Bayesian coalescent inference of past population dynamics from molecular sequences. Molecular Biology and Evolution, 22, 1185–1192.

Drummond, A. J., Suchard, M. A., Xie, D., and Rambaut, A. (2012). Bayesian phylogenetics with BEAUti and the BEAST 1.7. Molecular Biology and Evolution, 29, 1969–1973.

Edwards, C. T. T., Holmes, E. C., Pybus, O. G., Wilson, D. J., Viscidi, R. P., Abrams, E. J., Phillips, R. E., and Drummond, A. J. (2006). Evolution of the human immunodeficiency virus envelope gene is dominated by purifying selection. Genetics, 174, 1441–1453.

Ethier, S. N. and Griffiths, R. C. (1990). On the two-locus sampling distribution. Journal of Mathematical Biology, 29, 131–159.

Ewing, G., Nicholls, G., and Rodrigo, A. (2004). Using temporally spaced sequences to simultaneously estimate migration rates, mutation rate and population sizes in measurably evolving populations. Genetics, 168, 2407–2420.

Fan, J., Negroni, M., and Robertson, D. L. (2007). The distribution of HIV-1 recombination breakpoints. Infection, Genetics and Evolution, 7, 717–723.

Fearnhead, P. (2008). Computational methods for complex stochastic systems: a review of some alternatives to MCMC. Statistics and Computing, 18, 151–171.

Fearnhead, P. and Donnelly, P. (2001). Estimating recombination rates from population genetic data. Genetics, 159, 1299–1318.

Gall, A., Ferns, B., Morris, C., Watson, S., Cotten, M., Robinson, M., Berry, N., Pillay, D., and Kellam, P. (2012). Universal amplification, next-generation sequencing, and assembly of HIV-1 genomes. Journal of Clinical Microbiology, 50, 3838–3844.

Gall, A., Kaye, S., Hué, S., Bonsall, D., Rance, R., Baillie, G. J., Fidler, S. J., Weber, J. N., McClure, M. O., Kellam, P., and the SPARTAC Trial Investigators (2013). Restriction of V3 region sequence divergence in the HIV-1 envelope gene during antiretroviral treatment in a cohort of recent seroconverters. Retrovirology, 10, 8.

Grenfell, B. T., Pybus, O. G., Gog, J. R., Wood, J. L. N., Daly, J. M., Mumford, J. A., and Holmes, E. C. (2004). Unifying the epidemiological and evolutionary dynamics of pathogens. Science, 303, 327–332.

Griffiths, R. C. and Marjoram, P. (1996). Ancestral inference from samples of DNA sequences with recombination. Journal of Computational Biology, 3, 479–502.

Griffiths, R. C. and Tavar´e, S. (1994a). Sampling theory for neutral alleles in a varying environment. Philosophical Transactions of the Royal Society B, 344, 403–410.

Griffiths, R. C. and Tavar´e, S. (1994b). Simulating probability distributions in the coalescent. Theoretical Population Biology, 46, 131–159.

Griffiths, R. C., Jenkins, P. A., and Song, Y. S. (2008). Importance sampling and the two-locus model with subdivided population structure. Advances in Applied Probability, 40, 473–500.

Hein, J., Schierup, M. H., and Wiuf, C. Gene genealogies, variation and evolution. Oxford University Press, 2005.

Henn, M. R., Boutwell, C. L., Charlebois, P., Lennon, N. J., Power, K. A., Macalalad, A. R., Berlin, A. M., Malboeuf, C. M., Ryan, E. M., Gnerre, S., Zody, M. C., Erlich, R. L., Green, L. M., Berical, A., Wang, Y., Casali, M., Streeck, H., Bloom, A. K., Dudek, T., Tully, D., Newman, R., Axten, K. L., Gladden, A. D., Battis, L., Kemper, M., Zeng, Q., Shea, T. P., Gujja, S., Zedlack, C., Gasser, O., Brander, C., Hess, C., Günthard, H. F., Brumme, Z. L., Brumme, C. J., Bazner, S., Rychert, J., Tinsley, J. P., Mayer, K. H., Rosenberg, E., Pereyra, F., Levin, J. Z., Young, S. K., Jessen, H., Altfeld, M., Birren, B. W., Walker, B. D., and Allen, T. M. (2012). Whole genome deep sequencing of HIV-1 reveals the impact of early minor variants upon immune recognition during acute infection. PLoS Pathogens, 8, e1002529.

Jasra, A., De Iorio, M., and Chadeau-Hyam, M. (2011). The time machine: a simulation approach for stochastic trees. Proceedings of the Royal Society A: Mathematical, Physical and Engineering Sciences, 467, 2350–2368.

Jenkins, P. A. and Griffiths, R. C. (2011). Inference from samples of DNA sequences using a two-locus model. Journal of Computational Biology, 18, 109–127.

Jenkins, P. A. and Song, Y. S. (2009). Closed-form two-locus sampling distributions: accuracy and universality. Genetics, 183, 1087–1103.

Jukes, T. H. and Cantor, C. R. Evolution of protein molecules. In Mammalian protein metabolism, volume III, pages 21–132. Academic Press, New York, 1969.

Katoh, K., Misawa, K. K. Kuma, K., and Miyata, T. (2002). MAFFT: a novel method for rapid multiple sequence alignment based on fast Fourier transform. Nucleic Acids Research, 30, 3059–3066.

Kellam, P. and Larder, B. A. (1995). Retroviral recombination can lead to linkage of reverse transcriptase mutations that confer increased zidovudine resistance. Journal of Virology, 69, 669–674.

Kuhner, M. K., Yamato, J., and Felsenstein, J. (2000). Maximum likelihood estimation of recombination rates from population data. Genetics, 156, 1393–1401.

Larribe, F. and Fearnhead, P. (2011). On composite likelihoods in statistical genetics. Statistica Sinica, 21, 43–69.

Leblois, R., Pudlo, P., Néron, J., Bertaux, F., Beeravolu, C. R., Vitalis, R., and Rousset, F. (2014). Maximum likelihood inference of population size contractions from microsatellite data. Molecular Biology and Evolution, 31, 2805–2823.

Lee, A., Yau, C., Giles, M. B., Doucet, A., and Holmes, C. C. (2010). On the utility of graphics cards to perform massively parallel simulation of advanced Monte Carlo methods. Journal of Computational and Graphical Statistics, 19, 769–789.

Lemey, P., Rambaut, A., and Pybus, O. G. (2006). HIV evolutionary dynamics within and among hosts. AIDS reviews, 8, 125–40.

Lemey, P., Pond, S. L. K., Drummond, A. J., Pybus, O. G., Shapiro, B., Barroso, H., Taveira, N., and Rambaut, A. (2007). Synonymous substitution rates predict HIV disease progression as a result of underlying replication dynamics. PLoS Computational Biology, 3, e29.

Li, H. and Durbin, R. (2010). Fast and accurate long-read alignment with Burrows-Wheeler transform. Bioinformatics, 26, 589–595.

McVean, G., Awadalla, P., and Fearnhead, P. (2002). A coalescent-based method for detecting and estimating recombination from gene sequences. Genetics, 160, 1231–1241.

McVean, G. A. T., Myers, S. R., Hunt, S., Deloukas, P., Bentley, D. R., and Donnelly, P. (2004). The fine-scale structure of recombination rate variation in the human genome. Science, 304, 581–584.

Minin, V. N., Bloomquist, E. W., and Suchard, M. A. (2008). Smooth skyride through a rough skyline: Bayesian coalescent-based inference of population dynamics. Molecular Biology and Evolution, 25, 1459–1471.

Neher, R. A. and Leitner, T. (2010). Recombination rate and selection strength in HIV intra-patient evolution. PLoS Computational Biology, 6, e10000660.

Nielsen, R. (1997). A likelihood approach to populations samples of microsatellite alleles. Genetics, 146, 711–716.

Nowak, M. and May, R. M. Virus dynamics: mathematical principles of immunology and virology: mathematical principles of immunology and virology. Oxford University Press, 2000.

Paul, J. S. and Song, Y. S. (2010). A principled approach to deriving approximate conditional sampling distributions in population genetics models with recombination. Genetics, 186, 321–338.

Paul, J. S., Steinrücken, M., and Song, Y. S. (2011). An accurate sequentially Markov conditional sampling distribution for the coalescent with recombination. Genetics, 187, 1115–1128.

Pennings, P. S., Kryazhimskiy, S., and Wakeley, J. (2014). Loss and recovery of genetic diversity in adapting populations of HIV. PLoS Genetics, 10, e1004000.

Poon, A. F. Y., Swenson, L. C., Bunnik, E. M., Edo-Matas, D., Schuitemaker, H., van’t Wout, A. B., and Harrigan, P. R. (2012). Reconstructing the dynamics of HIV evolution within hosts from serial deep sequence data. PLoS Computational Biology, 8, e1’53.

Pybus, O. G. and Rambaut, A. (2009). Evolutionary analysis of the dynamics of viral infectious disease. Nature Reviews Genetics, 10, 540–550.

Pybus, O. G., Rambaut, A., and Harvey, P. H. (2000). An integrated framework for the inference of viral population history from reconstructed genealogies. Genetics, 155, 1429–1437.

Rasmussen, M. D., Hubisz, M. J., Gronau, I., and Siepel, A. (2014). Genome-wide inference of ancestral recombination graphs. PLOS Genetics, 10, e1004342.

Rodrigo, A. G. and Felsenstein, J. Coalescent approaches to HIV population genetics. In Crandall, K. A., editor, The evolution of HIV, pages 233–272. Johns Hopkins University Press, Baltimore, 1999.

Ross, H. A. and Rodrigo, A. G. (2002). Immune-mediated positive selection drives human immunodeficiency virus type 1 molecular variation and predicts disease duration. Journal of Virology, 76, 11715–11720.

Rouzine, I. M., Coffin, J. M., and Weinberger, L. S. (2014). Fifteen years later: Hard and soft selection sweeps confirm a large population number for HIV in vivo. PLoS Genetics, 10, e1004179.

Rouzine, I. and Coffin, J. (1999). Linkage disequilibrium test implies a large effective population number for HIV in vivo. Proceedings of the National Academy of Sciences, 96, 10758–10763.

Shankarappa, R., Margolick, J. B., Gange, S. J., Rodrigo, A. G., Upchurch, D., Farzadegan, H., Gupta, P., Rinaldo, C. R., Learn, G. H., He, X. I., Huang, X. L., and Mullins, J. I. (1999). Consistent viral evolutionary changes associated with the progression of human immunodeficiency virus type 1 infection. Journal of Virology, 73, 10489–10502.

Shapiro, B., Rambaut, A., and Drummond, A. J. (2006). Choosing appropriate substitution models for the phylogenetic analysis of protein-coding sequences. Molecular Biology and Evolution, 23, 7–9.

Sheehan, S., Harris, K., and Song, Y. S. (2013). Estimating variable effective population sizes from multiple genomes: a Sequentially Markov Conditional Sampling Distribution approach. Genetics, 194, 647–662.

Shriner, D., Rodrigo, A. G., Nickle, D. C., and Mullins, J. I. (2004). Pervasive genomic recombination of HIV-1 in vivo. Genetics, 167, 1573–1583.

Stephens, M. and Donnelly, P. (2000). Inference in molecular population genetics. Journal of the Royal Statistical Society: Series B, 62, 605–655.

Wang, Y. and Rannala, B. (2008). Bayesian inference of fine-scale recombination rates using population genomic data. Philosophical Transactions of the Royal Society B, 363, 3921–3930.

Williamson, S. (2003). Adaptation in the env gene of HIV-1 and evolutionary theories of disease progression. Molecular Biology and Evolution, 20, 1318–1325.

Wilson, I. J., Weale, M. E., and Balding, D. J. (2003). Inferences from DNA data: population histories, evolutionary processes and forensic match probabilities. Journal of the Royal Statistical Society: Series A, 166, 155–201.

